# Polymorphisms in Human Cytomegalovirus gO Exert Epistatic Influences on Cell-Free and Cell-To-Cell Spread, and Antibody Neutralization on gH Epitopes

**DOI:** 10.1101/867234

**Authors:** Le Zhang Day, Cora Stegmann, Eric P. Schultz, Jean-Marc Lanchy, Qin Yu, Brent J. Ryckman

## Abstract

The human cytomegalovirus (HCMV) glycoproteins H and L (gH/gL) can be bound by either gO, or the UL128-131 proteins to form complexes that facilitate entry and spread and the complexes formed are important targets of neutralizing antibodies. Strains of HCMV vary considerably in the levels of gH/gL/gO and gH/gL/UL128-131 and this can impact infectivity and cell tropism. In this report, we investigated how natural interstrain variation in the amino acid sequence of gO influences the biology of HCMV. Heterologous gO recombinants were constructed in which 6 of the 8 alleles or genotypes (GT) of gO were analyzed in the backgrounds of strain TR and Merlin (ME). The levels of gH/gL complexes were not affected, but there were impacts on entry, spread and neutralization by anti-gH antibodies. AD169 (AD) gO (GT1a) drastically reduced cell-free infectivity of both strains on fibroblasts and epithelial cells. PHgO(GT2a) increased cell-free infectivity of TR in both cell types, but spread in fibroblasts was impaired. In contrast, spread of ME in both cell types was enhanced by Towne (TN) gO (GT4), despite similar cell-free infectivity. TR expressing TNgO(GT4) was resistant to neutralization by anti-gH antibodies AP86 and 14-4b, whereas ADgO(GT1a) conferred resistance to 14-4b, but enhanced neutralization by AP86. Conversely, ME expressing ADgO(GT1a) was more resistant to 14-4b. These results suggest; 1) mechanistically distinct roles for gH/gL/gO in cell-free and cell-to-cell spread, 2) gO isoforms can differentially shield the virus from neutralizing antibodies, and 3) effects of gO polymorphisms are epistatically dependent on other variable loci.

**IMPORTANCE:** Advances in HCMV population genetics have greatly outpaced understanding of the links between genetic diversity and phenotypic variation. Moreover, recombination between genotypes may shuffle variable loci into various combinations with unknown outcomes. UL74(gO) is an important determinant of HCMV infectivity, and one of the most diverse loci in the viral genome. By analyzing interstrain heterologous UL74(gO) recombinants, we show that gO diversity can have dramatic impacts on cell-free and cell-to-cell spread as well as on antibody neutralization and that the manifestation of these impacts can be subject to epistatic influences of the global genetic background. These results highlight the potential limitations of laboratory studies of HCMV biology that use single, isolated genotypes or strains.

## INTRODUCTION

Recent application of state-of-the-art genomics approaches have begun to uncover a greater and more complex genetic diversity of human cytomegalovirus (HCMV) than had been appreciated (1–8). Of the 165 canonical open reading frames (ORFs) in the 235 kbp HCMV genome, 21 show particularly high nucleotide diversity and are distributed throughout the otherwise highly conserved genome. Links between specific genotypes and observed phenotypes are not well understood and as a corollary outcome, the factors driving HCMV genetic diversity and evolution remain speculative. This is further complicated by recombination between genotypes that can shuffle the diverse loci into various combinations, and this may result in epistasis where the phenotypic manifestation of a specific genotype of one locus may be influenced by the specific genotypes of other loci. Thus, realizing the full potential of modern genomics approaches towards the design of new interventions, clinical assessments and predictions will require better mechanistic understanding of the links between genotypes and phenotypes.

The UL74 ORF codes for glycoprotein (g) O and is one of the aforementioned highly diverse loci of HCMV (9–12). Most phylogenetic groupings indicate 8 genotypes or alleles of gO that differ in 10-30% of amino acids, predominately near the N-terminus and in a short central region. These amino acid polymorphisms also affect predicted N-linked glycan sites. The evolutionary origins of gO genotype diversity are not understood. Studies that followed infected humans through latency-reactivation cycles over several years demonstrated remarkable stability in UL74(gO) sequences, arguing against the idea of selective pressure from a dynamically adapting host immune system as a driving force for gO diversity (11, 13). The functional significance of gO diversity has only recently been addressed and centers around its role as a subunit of the envelope glycoprotein complex gH/gL/gO, which is involved in the initiation of infection into different cell types.

The general model for herpesvirus entry involves fusion between the virion envelope and cell membranes mediated by the fusion protein gB and the regulatory protein gH/gL (14–16). The HCMV gH/gL can be unbound, or bound by gO or the set of UL128-131 proteins (17–20). How these gH/gL complexes participate to mediate infection is complicated and seems to depend on both the cell type and whether the infection is by cell-free virus or direct cell-to-cell spread. Efficient infection of all cultured cell types by cell-free HCMV is dependent on gH/gL/gO, whereas infection of select cell types including epithelial and endothelial cells additionally requires gH/gL/UL128-131 (21–26). Experiments involving HCMV mutants lacking either gO or UL128-131 suggested that cell-to-cell spread in fibroblast cultures can be mediated by either gH/gL/gO or gH/gL/UL128-131, whereas in endothelial and epithelial cells gH/gL/UL128-131 is required, and it has remained unclear whether gH/gL/gO plays any role (23, 25, 27, 28). While it is clear that gH/gL/gO can bind to the cell surface protein PDGFRα via gO, and that gH/gL/UL128-131 can bind NRP2 and OR14I1 via UL128-131, the specific function(s) of these receptor engagements is unclear, but may include virion attachment, regulation of gB fusion activity, or activation of signal transduction pathways (29–31). In the case of gH/gL/gO, binding to PDGFRα activates signaling pathways, but these are not required for entry (28, 30, 32). Stegmann et al. showed that binding of a gO null HCMV to fibroblasts and endothelial cells was impaired, yet it is unclear whether this was due to lack of PDGFRα engagement. (33). Finally, Wu et al. reported coimmunoprecipitation of gB with gH/gL/gO and PDGFRα, consistent with a role for the gH/gL/gO-PDGFRα interaction in promoting gB fusion activity (32). However, unbound gH/gL has been shown to mediate cell-cell fusion and has also been found in stable complex with gB in extracts of infected cells and extracellular virions (20, 34). Thus, although many of the key factors in HCMV entry and cell-to-cell spread have been identified, their interplay in the various entry pathways is unclear. Moreover, the influence of gO diversity remains a mystery.

The gH/gL complexes have been extensively studied as potential vaccine candidates and neutralizing antibodies have been described that react with epitopes on gH/gL, on UL128-131 and on gO (35–43). Anti-UL128-131 antibodies neutralize with high potency, but only on cell types for which gH/gL/UL128-131 is required for entry; e.g., epithelial cells. In contrast, antibodies that react with epitopes on gH/gL tend to neutralize virus on both fibroblasts and epithelial cells, but are far less potent on fibroblasts, where only gH/gL/gO is needed for entry. One explanation for these observations is that gO, with its extensive N-linked glycan decorations presents more steric hindrance to antibodies accessing the underlying gH/gL epitopes than do the UL128-131 proteins. Similar effects of glycans in shielding neutralizing epitopes have been described for HIV env, and for HCMV gN (44) (45). In support of this hypothesis for gO, Jiang et al. showed that focal spread of a gO null HCMV in fibroblasts was more sensitive to anti-gH antibodies (46). Recently, Cui et al. described antibodies that reacted to a linear epitope on gH that exhibited strain-selective neutralization that could not be explained by polymorphisms within the gH epitope (47). One possible explanation was that gO polymorphisms between the strains imposed differential steric hindrances on these antibodies.

In this study we utilized a set of HCMV BAC-clones that represent the range of phenotypic diversity in terms of gH/gL complexes. HCMV TB40/e (TB), TR and Merlin (ME) differ dramatically in the amounts of gH/gL complexes in the virion envelope and their infectivity on fibroblasts and epithelial cells. Extracellular virions of TB and TR contain gH/gL predominately in the form of gH/gL/gO and are far more infectious on both fibroblasts and epithelial cells than ME, which contains overall lower amounts of gH/gL, predominately as gH/gL/UL128-131 (9, 26). Each of these strains encodes a different representative of the 8 gO genotypes. In a previous report, we demonstrated that variation in the UL74(gO) ORF was not responsible for the observed differences between TR and ME. (48). Rather, it was shown that the amounts of gH/gL/gO in ME and TR virions were influenced by different steady-state levels of gO present during progeny assembly. Kalser et al. showed that replacing the gO of TB with that of Towne (TN) also did not affect the levels of gH/gL complexes but may have enhanced the ability of TB to spread in epithelial cell cultures (49). Here, we have generated a set of heterologous gO recombinants to include 6 of the 8 genotypes in the genetic backgrounds of the gH/gL/gO-rich strain TR and the gH/gL/UL128-131-rich ME to analyze how the differences in gO sequence influence HCMV biology. The results demonstrate that gO variation can have dramatic effects on cell-free entry, cell-to-cell spread and the neutralization by anti-gH antibodies. In some cases opposite influences were observed for a given gO genotype in the different backgrounds of TR and ME, indicating epistasis with other genetic differences between these strains.

## RESULTS

### Influences of gO polymorphisms on cell-free infectivity and tropism can be dependent on the background strain

To examine the effects of gO polymorphism, a set of recombinant viruses was constructed in which the endogenous UL74(gO) ORFs of strain TR and ME were replaced with the UL74(gO) ORFs from 5 other strains. BAC-cloned strains TR and ME were chosen as the backgrounds for these studies since they represent gH/gL/gO-rich and gH/gL/UL128-131-rich strains respectively (9, 26, 49). Additionally, ME is restricted to a cell-to-cell mode of spread in culture, whereas TR is capable of both cell-free and cell-to-cell modes of spread (23, 50, 51). The intended changes to UL74(gO) in each recombinant BAC were verified by sequencing the UL74 ORF and the flanking regions used for BAC recombineering. However, it was recently reported that HCMV BAC-clones can sustain various genetic deletions, and rearrangements, and mutations during rescue in fibroblasts or epithelial cells, resulting in mixed genotype populations (52). To ensure that phenotypes characterized were the associated with the intended changes to UL74(gO) and not to other genetic changes sustained during BAC rescue in fibroblasts, all analyses were performed on at least three independently BAC-rescued viral stocks.

As a basis for interpretation of the later biological comparisons among recombinants, the levels of gH/gL complexes incorporated into the virion envelope were analyzed by immunoblot as previously described (9, 26). As in the previous reports, TR contained predominantly gH/gL/gO, whereas ME contained mostly gH/gL/UL128-131 (Fig 1, compare lane 1 in panels A and B). Propagation of ME under conditions of UL131 transcriptional repression (denoted “Merlin-T” (MT) as described (26, 51)), resulted in more gH/gL/gO and less gH/gL/UL128-131 (Fig. 1C, lane 1). Some minor differences in the amounts of total gL, gH/gL/gO, and gH/gL/UL128-131 were observed for some of the heterologous gO recombinants relative to their parental strains. However, band density analyses showed that all apparent differences were less than 3-fold and few reached statistical significance when compared across multiple experiments, likely reflecting the limitations of immunoblot as a precise quantitative method, as well as stock-to-stock variability in glycoprotein composition (Table 1). Thus, consistent with our previous report, differences between strains TR and ME in the abundance of gH/gL complexes are predominately influenced by genetic background differences outside the UL74(gO) ORF (48).

**Figure 1.**
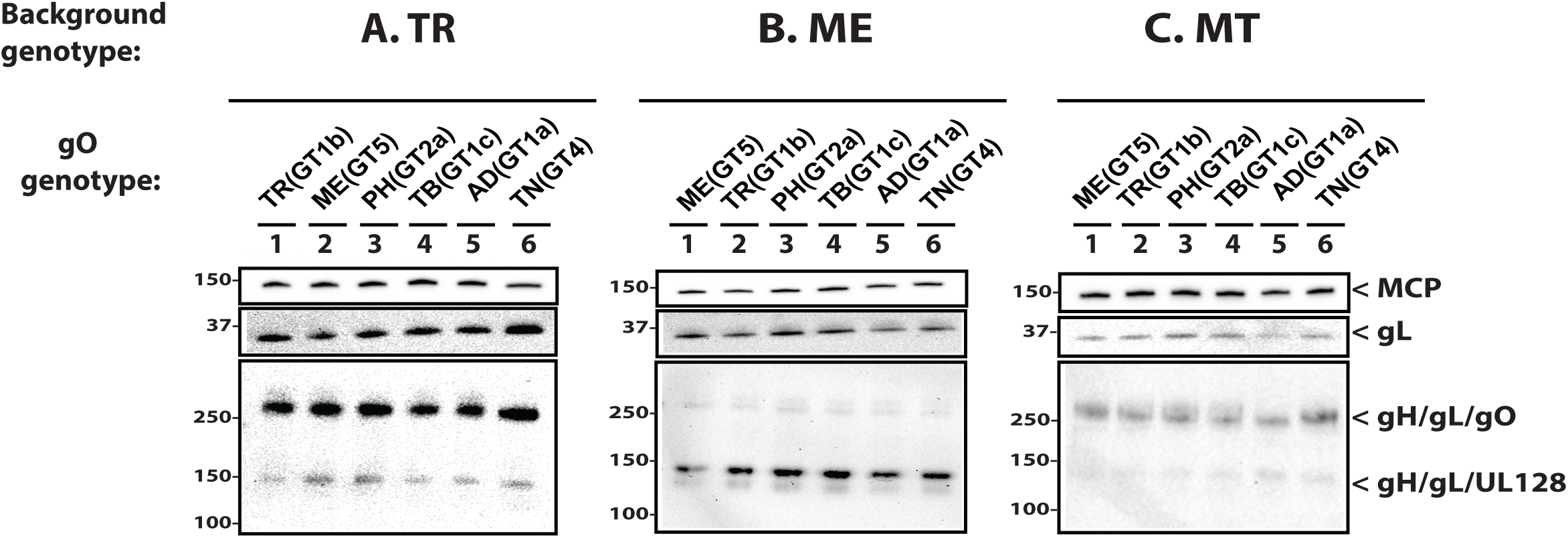
Immunoblot analysis of gH/gL complexes in parental and heterologous gO recombinant HCMV. Equal number of cell-free virions (as determined by qPCR) of HCMV TR (A), ME (B), or MT (C) or the corresponding heterologous gO recombinants were separated by reducing (upper two panels) or non-reducing (bottom panel) SDS-PAGE, and analyzed by immunoblot with antibodies specific for major capsid protein (MCP) or gL. Blots shown are representative of three independent experiments. Molecular mass markers (kDa) indicated on each panel.

**Table 1.** Immunoblot band density analyses of parental and heterologous gO recombinants.

| Genotype Background | Virion Protein(s) Analyzed |  |  |  |  |  |  |  |
| --- | --- | --- | --- | --- | --- | --- | --- | --- |
| TR | MCP |  | gL |  | gH/gL/gO |  | gH/gL/UL128 |  |
| <u>gO genotype</u> | <u>Fold<sup>b</sup></u> | <u>ANOVA<sup>c</sup></u> | <u>Fold</u> | <u>p-value</u> | <u>Fold</u> | <u>ANOVA</u> | <u>Fold</u> | <u>ANOVA</u> |
| TR(GT1b) | - | - | - | - | - | - | - | - |
| MEgO(GT5) | 1.1 | ns | 0.6 | ns | 1.4 | ns | 2.0 | ns |
| PHgO(GT2a) | 1.1 | ns | 0.9 | ns | 1.8 | ns | 2.3 | * |
| TBgO (GT1c) | 1.2 | ns | 0.8 | ns | 0.9 | ns | 0.9 | ns |
| ADgO (GT1a) | 1.1 | ns | 0.9 | ns | 0.9 | ns | 1.0 | ns |
| TNgO (GT4) | 1.1 | ns | 2.0 | ns | 2.7 | ns | 2.1 | ns |
| ME | MCP |  | gL |  | gH/gL/gO |  | gH/gL/UL128 |  |
| <u>gO genotype</u> | <u>Fold</u> | <u>ANOVA</u> | <u>Fold</u> | <u>ANOVA</u> | <u>Fold</u> | <u>ANOVA</u> | <u>Fold</u> | <u>ANOVA</u> |
| MEgO(GT5) | - | - | - | - | - | - | - | - |
| TR(GT1b) | 0.9 | ns | 0.8 | ns | 0.9 | ns | 1.1 | ns |
| PHgO(GT2a) | 1.1 | ns | 1.1 | ns | 1.4 | ns | 1.4 | ns |
| TBgO (GT1c) | 1.3 | ns | 1.2 | ns | 1.0 | ns | 1.4 | ns |
| ADgO (GT1a) | 1.0 | ns | 0.7 | ns | 0.9 | ns | 1.1 | ns |
| TNgO (GT4) | 1.1 | ns | 0.8 | ns | 0.9 | ns | 1.4 | ns |
| MT | MCP |  | gL |  | gH/gL/gO |  | gH/gL/UL128 |  |
| <u>gO genotype</u> | <u>Fold</u> | <u>ANOVA</u> | <u>Fold</u> | <u>ANOVA</u> | <u>Fold</u> | <u>ANOVA</u> | <u>Fold</u> | <u>ANOVA</u> |
| MEgO(GT5) | - | - | - | - | - | - | - | - |
| TR(GT1b) | 1.1 | ns | 1.2 | ns | 0.7 | ns | 0.9 | ns |
| PHgO(GT2a) | 1.1 | ns | 1.6 | ns | 1.4 | ns | 1.1 | ns |
| TBgO (GT1c) | 1.1 | ns | 1.3 | ns | 1.1 | ns | 1.6 | ns |
| ADgO (GT1a) | 0.8 | ns | 0.5 | ns | 0.6 | ns | 1.7 | ns |
| TNgO (GT4) | 0.9 | ns | 0.7 | ns | 1.4 | ns | 1.8 | ns |

b. Mean fold difference of chemiluminescent band densities obtained for each recombinant compared to the parental TR in three independent experiments.
c. One-way ANOVA with Dunnett’s multiple comparisons test comparing each recombinant to the parental in three independent experiments. (\*) $p \leq 0.05$ , (ns) not significant.

While gH/gL/gO is clearly important for entry into both fibroblasts and epithelial cells, the mechanisms are likely different since 1) fibroblasts clearly express the gH/gL/gO receptor PDGFRα on their surface, whereas ARPE19 epithelial cells express little or none of this protein (28, 30, 32, 53), and 2) entry into epithelial cells requires gH/gL/UL128-131 in addition to gH/gL/gO (23, 24, 26). Thus, it was possible that gO polymorphisms would differentially affect replication in these two cell types. To address this, fibroblast-to-epithelial tropism ratios were determined for each parental strain and gO recombinant by inoculating cultures of fibroblasts and epithelial cells in parallel with equivalent amounts of cell-free virus stocks. The number of infected cells in each culture was then determined by flow cytometry using GFP expressed from the virus genome. Figure 2 shows the results of these experiments as the fold preference for either cell type as a ratio, where “1” indicates equal infection of both cell types. Stocks of the parental TR were approximately 20-fold more infectious on fibroblasts than on epithelial cells (Fig 2A). Preference towards fibroblasts was greater for TR-recombinants expressing MEgO(GT5), PHgO(GT2b), and TBgO(GT1c). In contrast, tropism ratios of TR-recombinants expressing ADgO(GT1a) and TNgO(GT4) were closer to 1, indicating more equal infection of both cell types. Parental ME and all of the ME-based gO recombinants had tropism ratios within the range of 6 in favor of fibroblasts to 3 in favor of epithelial cells. Several of these viruses had variability between replicate stocks where some had slight fibroblasts preference and others slight epithelial preference (Fig 2B). Propagation of the ME-based viruses as MT greatly increased the preference towards fibroblasts infection for all recombinants to a range of 30-300 fold (Fig 2B). These results suggested that for the more gH/gL/gO-rich TR and MT, gO polymorphisms may differentially influence the infection of fibroblasts and epithelial cells, shifting the apparent relative tropism. However, such influences were less pronounced for ME, consistent with the low abundance of gH/gL/gO expressed by this virus.

**Figure 2.**
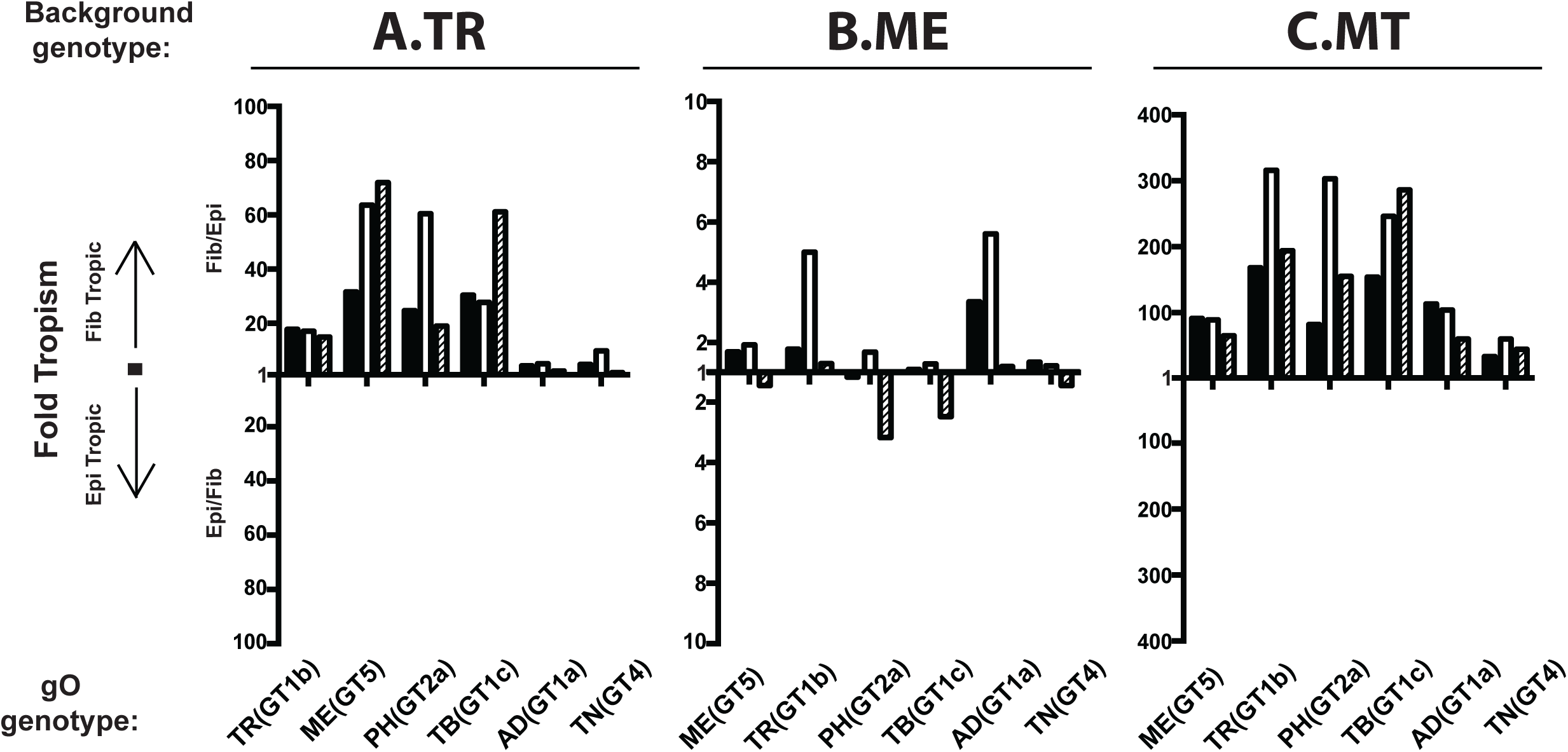
Relative fibroblast and epithelial cell tropism of parental and heterologous gO recombinant. HCMV. Cell-free stocks of HCMV TR (A), ME (B), or MT (C) or the corresponding heterologous gO recombinants were serially diluted, and side-by-side cultures of nHDF fibroblasts and ARPE19 epithelial cells were inoculated with equal volumes of the dilutions. The number of infected cells was determined by flow cytometry for GFP at 2 days post infection. Ratios greater than or equal to 1 of the number of each cell type infected (fib/epi or epi/fib) are plotted for each of three independent sets of virus stocks (black, open and striped bars).

It was not clear if the observed differences in tropism ratios were due to enhanced infection of one cell type, reduced infection of the other cell type or a mixture of both. To address this, specific infectivity (ratio of the number of virions to the number of infectious units) was determined for each parental and recombinant on both fibroblasts and epithelial cells. Multiple independent supernatant stocks of each recombinant were analyzed by qPCR for encapsidated viral genomes and infectious titers on both cell types were determined by flow cytometry quantification of GFP-positive cells (Fig 3). For the TR-based viruses on fibroblasts, MEgO(GT5), TBgO(GT1c), and TNgO(GT4) each resulted in moderately enhanced infectivity (2 to 10-fold fewer genomes/IU) compared to the parental TR, and PHgO(GT2a) enhanced infectivity 30-fold. In contrast, ADgO(GT1a) dropped TR infectivity below the detection limit of the flow cytometry-based assay (Fig 3A, top panel). In our previous report, expression of MEgO in the TR background did not appear to affect infectivity on fibroblasts (48). This discrepancy was likely due to the more sensitive flow cytometry readout used in the current studies as compared to the plaque assay readout used previously. The infectivity of parental TR on epithelial cells was about 20-fold lower than on fibroblasts (i.e., 20-fold higher genomes/IU), but the relative effect of each heterologous gO was similar to that observed on fibroblasts (Fig 3A, bottom panel). Thus, some of the gO changes had dramatic effects on the infectivity of TR. Although these effects were manifest on both cell types, they were more pronounced on fibroblasts and this explains the observed differences in fibroblast preferences reported in Figure 2A.

**Figure 3.**
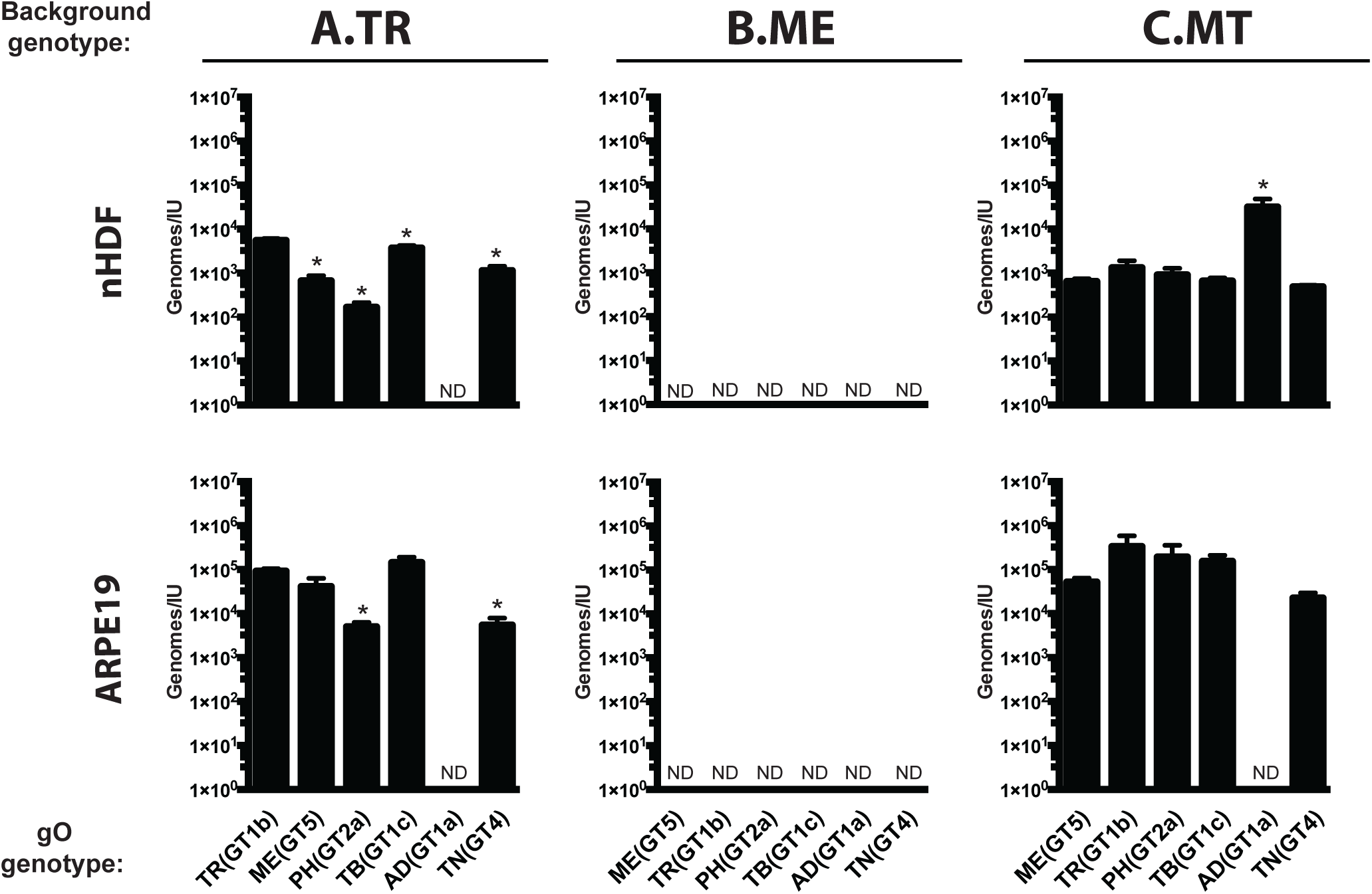
Specific infectivity of parental and heterologous gO recombinant HCMV. Extracellular HCMV stocks of HCMV TR (A), ME (B), or MT (C) or the corresponding heterologous gO recombinants were quantified by qPCR for viral genomes, and infectious units (IU) were determined by flow cytometry quantification of GFP-expressing nHDF fibroblasts or ARPE-19 epithelial cells, 2 days post infection. Average genomes/IU of 3 independent set of virus stock are plotted, with error bars representing standard deviations. Undetectable levels of infectivity indicated by ND (not determined). Asterisks (*) denote p-values ≤ 0.05; one-way ANOVA with Dunnett’s multiple comparisons test comparing each recombinant to the parental in three independent experiments.

The infectivity of cell-free ME virions on both cell types was below the detection limit of the flow cytometry-based assay and none of the changes to gO rescued infectivity (Fig 3B). These results indicated that the cell-free virions of all of the ME-based viruses were virtually non-infectious. When propagated as MT, infectivity on both cell types was improved to levels comparable to TR and this was consistent with our previous results (Fig 2C) (26, 48). The only significant effect of gO changes on MT was ADgO(GT1a), which reduced infectivity on both cell types,. Thus, as in the TR background, some changes to gO influenced infectivity of MT and this was disproportionally manifest on fibroblasts compared epithelial cells, but the overall preference of all of the MT-based viruses was strongly in favor of fibroblasts. In contrast, gO changes had little effect on the infectivity or tropism of ME-based viruses.

It has been reported that gO-null HCMV are impaired for attachment to cells and that soluble gH/gL/gO can block HCMV attachment (33, 54). Thus, it was possible that the observed changes to cell-free infectivity due to gO polymorphisms were related to a role for gO in attachment. To test this hypothesis, each heterologous gO recombinant was compared to the corresponding parental strain by applying cell-free virus stocks to fibroblast or epithelial cell cultures for approximately 20 min, washing away the unbound virus and then counting the numbers of cell-associated virions by immunofluorescence staining of the capsid-associated tegument protein pp150 (33) (Fig 4 and Tables 2 and 3). Given the short incubation time, high concentrations of input viruses were used to, and these inputs were equal for each set of parental and heterologous gO recombinants within the constraints of the stock concentrations. Higher inputs were required for ME to obtain detectable numbers of bound virus, consistent with the low amounts of gH/gL/gO in these virions. The average number of cell-associated virions per cell varied considerable between experiments, likely reflecting the complex parameters expected to influence virus attachment including stock concentration, cell state and variability in the incubation time between experiments. In some cases, a given recombinant was significantly different from parental in only one or two of the three experiments. It was concluded that these specific gO isoforms did not affect binding or attachment of HCMV to cells. However, binding of TR_TNgO(GT4) and MT_ADgO(GT1a) were each significantly lower than their respective parental viruses in all three experiments on both fibroblasts and epithelial cells. While it was possible that the reduced binding of MT_ADgO(GT1a) was due in part to the slightly lower amounts of gH/gL/gO (Fig 1C and Table 1), the reduced binding of TR_TNgO(GT4) could not be similarly explained since this virus had slightly more gH/gL/gO than the parental TR (Fig. 1A, Table 1). Moreover, reduced binding may help explain the lower infectivity of MT_ADgO(GT1a) (Fig 3C), but the poor infectivity of TR_ADgO(GT1a) could not be explained by poor binding, and the reduced binding of TR_TNgO(GT4) did not result in reduction of infectivity (Fig 3A).

**Figure 4.**
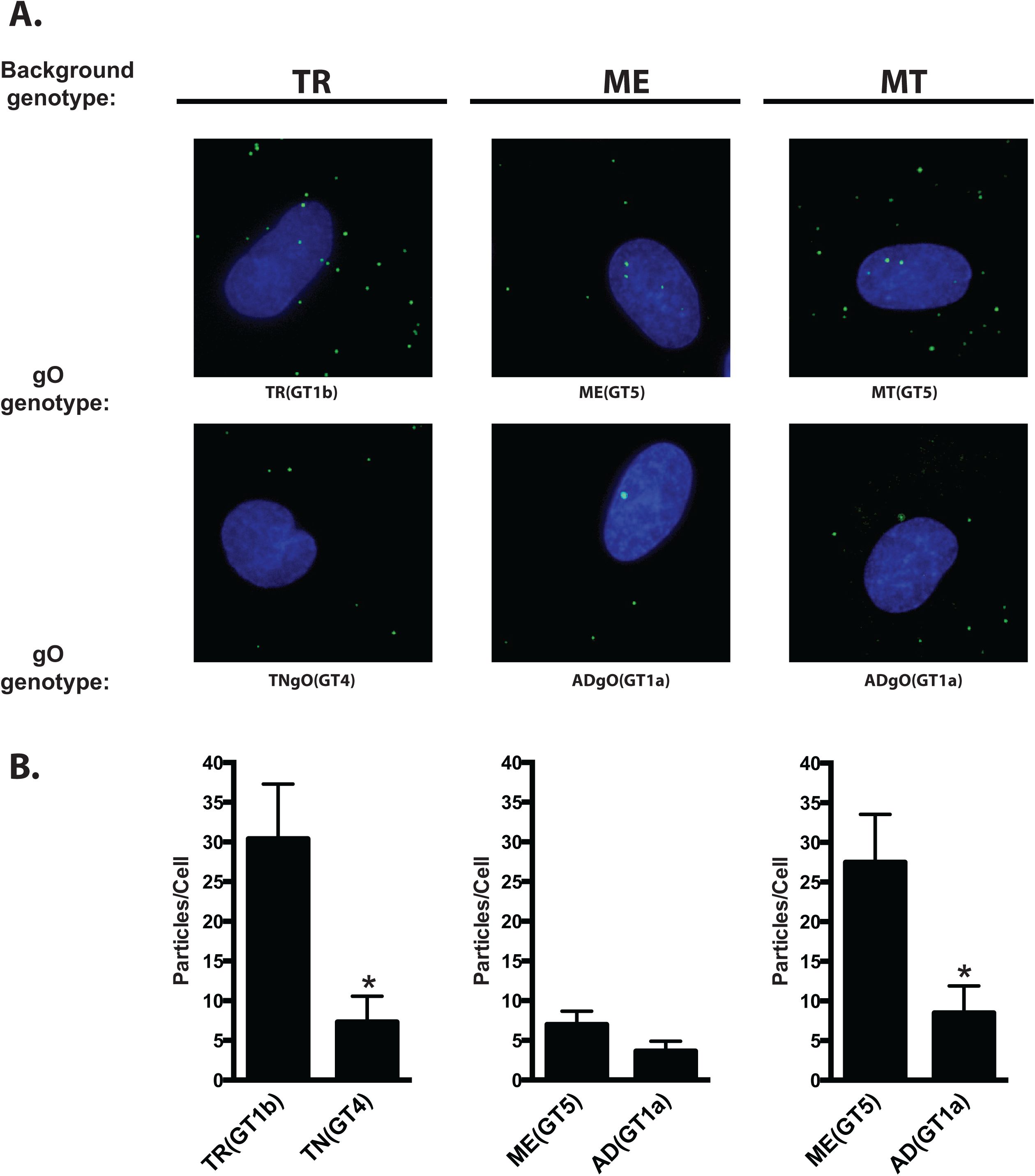
Binding of parental and heterologous gO recombinant HCMV to fibroblasts. Extracellular HCMV TR, ME, MT or the corresponding heterologous gO recombinants were applied to nHDF for 20 min. Multiplicities (genomes/cell) were: TR-background viruses (1 x 10^4^), ME-background viruses (5 x 10^4^), MT-background viruses (1 x 10^4^). After washing away unbound virus, cultures were fixed and permeabilized with acetone and cell-associated virus particles were detected by immunofluorescence using antibodies specific for the capsid-associated tegument protein, pp150. Cells were visualized by staining nuclei with DAPI. (A) Representative fields of parental TR, ME, MT and heterologous gO recombinants that consistently reduced binding in 3 independent experiments (Table 2). (B) Mean particles per cell for representative experiments. Error bars represent the standard deviation. Asterisks (*) denote p-values ≤ 0.05; one-way ANOVA with Dunnett’s multiple comparisons test comparing each recombinant to the parental.

**Table 2.**
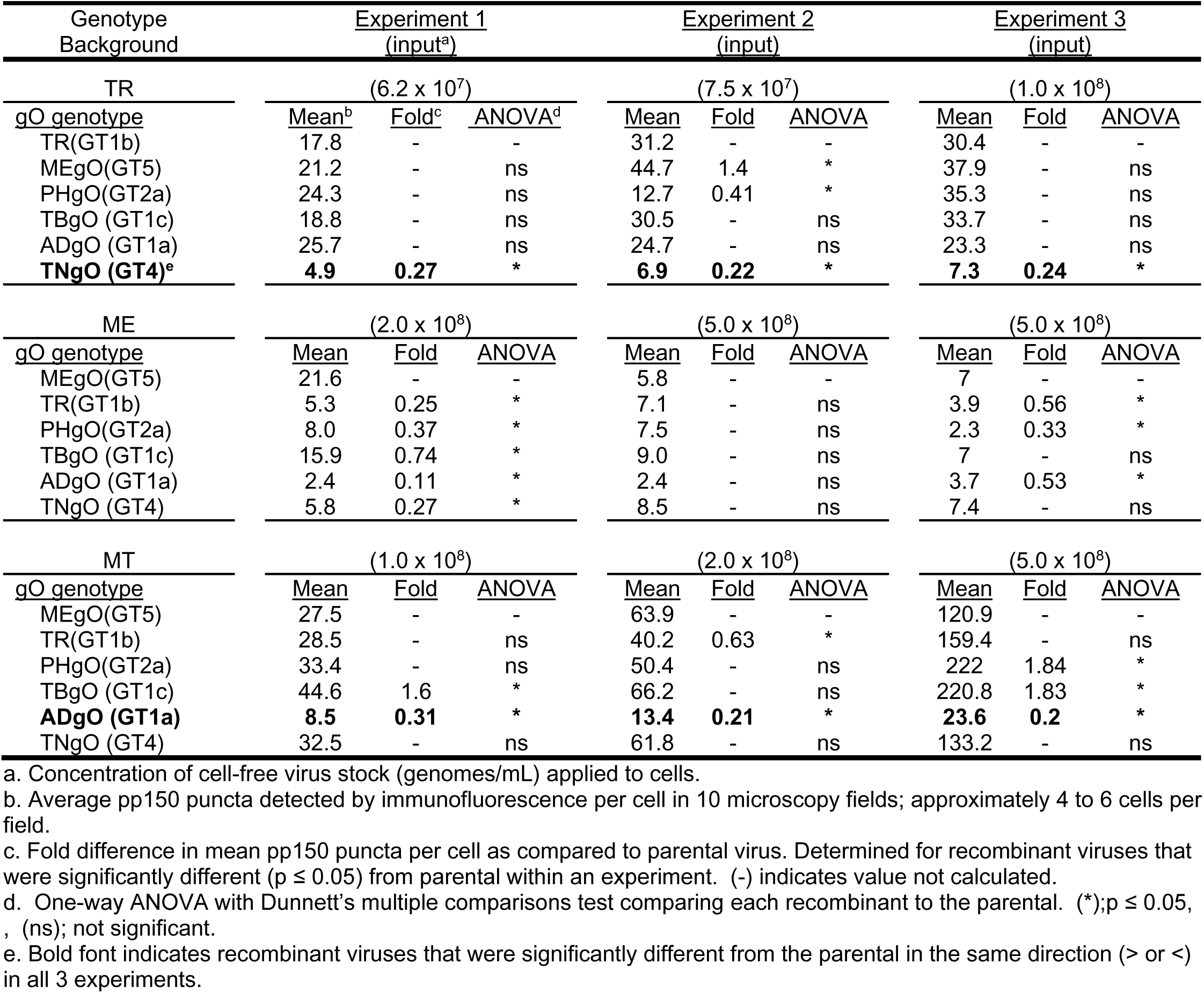
Binding of parental and heterologous gO recombinant HCMV to fibroblasts.

| Genotype Background | Experiment 1<br>(input<sup>a</sup>) |  |  | Experiment 2<br>(input) |  |  | Experiment 3<br>(input) |  |  |
| --- | --- | --- | --- | --- | --- | --- | --- | --- | --- |
| TR | (6.2 x 10<sup>7</sup>) |  |  | (7.5 x 10<sup>7</sup>) |  |  | (1.0 x 10<sup>8</sup>) |  |  |
| <u>gO genotype</u> | <u>Mean<sup>b</sup></u> | <u>Fold<sup>c</sup></u> | <u>ANOVA<sup>d</sup></u> | <u>Mean</u> | <u>Fold</u> | <u>ANOVA</u> | <u>Mean</u> | <u>Fold</u> | <u>ANOVA</u> |
| TR(GT1b) | 17.8 | - | - | 31.2 | - | - | 30.4 | - | - |
| MEgO(GT5) | 21.2 | - | ns | 44.7 | 1.4 | * | 37.9 | - | ns |
| PHgO(GT2a) | 24.3 | - | ns | 12.7 | 0.41 | * | 35.3 | - | ns |
| TBgO (GT1c) | 18.8 | - | ns | 30.5 | - | ns | 33.7 | - | ns |
| ADgO (GT1a) | 25.7 | - | ns | 24.7 | - | ns | 23.3 | - | ns |
| **TNgO (GT4)<sup>e</sup>** | **4.9** | **0.27** | * | **6.9** | **0.22** | * | **7.3** | **0.24** | * |
| ME | (2.0 x 10<sup>8</sup>) |  |  | (5.0 x 10<sup>8</sup>) |  |  | (5.0 x 10<sup>8</sup>) |  |  |
| <u>gO genotype</u> | <u>Mean</u> | <u>Fold</u> | <u>ANOVA</u> | <u>Mean</u> | <u>Fold</u> | <u>ANOVA</u> | <u>Mean</u> | <u>Fold</u> | <u>ANOVA</u> |
| MEgO(GT5) | 21.6 | - | - | 5.8 | - | - | 7 | - | - |
| TR(GT1b) | 5.3 | 0.25 | * | 7.1 | - | ns | 3.9 | 0.56 | * |
| PHgO(GT2a) | 8.0 | 0.37 | * | 7.5 | - | ns | 2.3 | 0.33 | * |
| TBgO (GT1c) | 15.9 | 0.74 | * | 9.0 | - | ns | 7 | - | ns |
| ADgO (GT1a) | 2.4 | 0.11 | * | 2.4 | - | ns | 3.7 | 0.53 | * |
| TNgO (GT4) | 5.8 | 0.27 | * | 8.5 | - | ns | 7.4 | - | ns |
| MT | (1.0 x 10<sup>8</sup>) |  |  | (2.0 x 10<sup>8</sup>) |  |  | (5.0 x 10<sup>8</sup>) |  |  |
| <u>gO genotype</u> | <u>Mean</u> | <u>Fold</u> | <u>ANOVA</u> | <u>Mean</u> | <u>Fold</u> | <u>ANOVA</u> | <u>Mean</u> | <u>Fold</u> | <u>ANOVA</u> |
| MEgO(GT5) | 27.5 | - | - | 63.9 | - | - | 120.9 | - | - |
| TR(GT1b) | 28.5 | - | ns | 40.2 | 0.63 | * | 159.4 | - | ns |
| PHgO(GT2a) | 33.4 | - | ns | 50.4 | - | ns | 222 | 1.84 | * |
| TBgO (GT1c) | 44.6 | 1.6 | * | 66.2 | - | ns | 220.8 | 1.83 | * |
| **ADgO (GT1a)** | **8.5** | **0.31** | * | **13.4** | **0.21** | * | **23.6** | **0.2** | * |
| TNgO (GT4) | 32.5 | - | ns | 61.8 | - | ns | 133.2 | - | ns |

a. Concentration of cell-free virus stock (genomes/mL) applied to cells.
b. Average pp150 puncta detected by immunofluorescence per cell in 10 microscopy fields; approximately 4 to 6 cells per field.
c. Fold difference in mean pp150 puncta per cell as compared to parental virus. Determined for recombinant viruses that were significantly different ( $p \leq 0.05$ ) from parental within an experiment. (-) indicates value not calculated.
d. One-way ANOVA with Dunnett's multiple comparisons test comparing each recombinant to the parental. (\*); $p \leq 0.05$ , (ns); not significant.
e. Bold font indicates recombinant viruses that were significantly different from the parental in the same direction (> or <) in all 3 experiments.

**Table 3.** Binding of parental and heterologous gO recombinant HCMV to epithelial cells.

| Genotype<br>Background | Experiment 1<br>(input<sup>a</sup>) |  |  | Experiment 2<br>(input) |  |  | Experiment 3<br>(input) |  |  |
| --- | --- | --- | --- | --- | --- | --- | --- | --- | --- |
| TR | (6.2 x 10<sup>7</sup>) |  |  | (7.5 x 10<sup>7</sup>) |  |  | (1.0 x 10<sup>8</sup>) |  |  |
| <u>gO genotype</u> | <u>Mean<sup>b</sup></u> | <u>Fold<sup>c</sup></u> | <u>ANOVA<sup>d</sup></u> | <u>Mean</u> | <u>Fold</u> | <u>ANOVA</u> | <u>Mean</u> | <u>Fold</u> | <u>ANOVA</u> |
| TR(GT1b) | 26.2 | - | - | 41.7 | - | - | 43.7 | - | - |
| MEgO(GT5) | 35.5 | 1.35 | * | 38.3 | - | ns | 56.8 | - | ns |
| PHgO(GT2a) | 33.4 | - | ns | 19.3 | 0.46 | * | 61 | 1.4 | * |
| TBgO (GT1c) | 24.1 | - | ns | 35.4 | - | ns | 58.7 | 1.34 | * |
| ADgO (GT1a) | 36.4 | 1.39 | * | 22.2 | 0.53 | * | 36 | - | ns |
| **TNgO (GT4)<sup>e</sup>** | **16.2** | **0.62** | * | **18.62** | **0.45** | * | **23.4** | **0.54** | * |
| ME | (2.0 x 10<sup>8</sup>) |  |  | (5.0 x 10<sup>8</sup>) |  |  | (5.0 x 10<sup>8</sup>) |  |  |
| <u>gO genotype</u> | <u>Mean</u> | <u>Fold</u> | <u>ANOVA</u> | <u>Mean</u> | <u>Fold</u> | <u>ANOVA</u> | <u>Mean</u> | <u>Fold</u> | <u>ANOVA</u> |
| MEgO(GT5) | 37.3 | - | - | 18 | - | - | 15 | - | - |
| TR(GT1b) | 17.7 | 0.47 | * | 24.9 | - | ns | 10.4 | 0.69 | * |
| PHgO(GT2a) | 22.3 | 0.6 | * | 23 | - | ns | 9.4 | 0.62 | * |
| TBgO (GT1c) | 34.1 | - | ns | 32.3 | 1.79 | * | 18.6 | - | ns |
| ADgO (GT1a) | 14.4 | 0.39 | * | 11.4 | - | ns | 10.8 | 0.72 | * |
| TNgO (GT4) | 24.4 | 0.65 | * | 25.9 | 1.44 | * | 14.3 | - | ns |
| MT | (1.0 x 10<sup>8</sup>) |  |  | (2.0 x 10<sup>8</sup>) |  |  | (5.0 x 10<sup>8</sup>) |  |  |
| <u>gO genotype</u> | <u>Mean</u> | <u>Fold</u> | <u>ANOVA</u> | <u>Mean</u> | <u>Fold</u> | <u>ANOVA</u> | <u>Mean</u> | <u>Fold</u> | <u>ANOVA</u> |
| MEgO(GT5) | 33.2 | - | - | 68 | - | - | 236.8 | - | - |
| TR(GT1b) | 35.3 | - | ns | 46.1 | 0.68 | * | 210.1 | - | ns |
| PHgO(GT2a) | 46.5 | - | ns | 78 | - | ns | 383.2 | 1.62 | * |
| TBgO (GT1c) | 63.4 | 1.91 | * | 69.6 | - | ns | 238.3 | - | ns |
| **ADgO (GT1a)** | **16.7** | **0.5** | * | **26.1** | **0.38** | * | **26.6** | **0.11** | * |
| TNgO (GT4) | 44.1 | - | ns | 48.1 | 0.71 | * | 150.9 | 0.64 | * |

a. Concentration of cell-free virus stock (genomes/mL) applied to cells.
b. Average pp150 puncta detected by immunofluorescence per cell in 10 microscopy fields; approximately 4 to 6 cells per field.
c. Fold difference in mean pp150 puncta per cell as compared to parental virus. Determined for recombinant viruses that were significantly different ( $p \leq 0.05$ ) from parental within an experiment. (-) indicates value not calculated.
d. One-way ANOVA with Dunnett's multiple comparisons test comparing each recombinant to the parental. (\*) $p \leq 0.05$ , (ns) not significant
e. Bold font indicates recombinant viruses that were significantly different from the parental in the same direction (> or <) in all 3 experiments.

In sum, these analyses indicated that; 1) gO polymorphisms can influence the cell-free infectivity of HCMV. In some cases this was independent of any effects on abundance of gH/gL/gO in the virion envelope or binding to cells (e.g. parental TR and TR recombinants harboring MEgO(GT5), TBgO(GT1c), and ADgO(GT1a), had dramatically different infectivity but comparable levels of gH/gL/gO and cell binding). 2) The influence of some gO isoforms was dependent on the background strain (e.g., PHgO(GT2a) enhanced TR infectivity but did not affect ME or MT and TNgO(GT4) reduced binding of TR but had no effect on binding of ME or MT). 3) While some heterologous gO recombinants had quantitatively different effects on infectivity on fibroblast compared to epithelial cells, these did not change the fundamental fibroblast preferences for either TR or MT. 4) Some of the heterologous gOs did appear to change relative tropism of ME. However, the relevance of tropism ratios for these viruses is questionable since the specific infectivity (genomes/IU) analyses suggested that all ME-based recombinants were noninfectious on either cell type. This was consistent with the highly cell-associated nature of ME (50, 51).

### Polymorphisms in gO can differentially influence the mechanisms of cell-free and cell-to-cell spread

The analyses described above focused on the cell-free infectivity of HCMV, as indicative of a cell-free mode of spread. Cell-to-cell spread mechanisms are likely important for HCMV, and while gH/gL complexes are clearly important for cell-to-cell spread, the mechanisms in these processes are poorly characterized in comparison to cell-free infection. Strains TR and ME are well-suited to compare the effects of gO polymorphisms on cell-free and cell-to-cell spread since ME is mostly restricted to cell-to-cell due to the poor infectivity of cell-free virions but can be allowed to also spread cell-free by propagation as MT, whereas TR can spread by both cell-free and cell-to-cell mechanisms (23, 26, 50, 51).

To compare spread among heterologous gO recombinants, replicate cultures were infected at low multiplicity, and at 12 dpi, foci morphology was documented by fluorescence microscopy and the increased number of infected cells was determined by flow cytometry. In fibroblasts cultures, parental TR and MT showed more diffuse foci compared to the tight, localized focal pattern of parental ME, consistent with the notion that TR and MT spread by both cell-free and cell-to-cell mechanisms whereas ME was restricted to cell-to-cell spread (Fig 5A). Quantitatively, spread by parental TR increased the numbers of infected cells 55-fold over 12 days, whereas spread of TR_MEgO(GT5) and TR_PHgO(GT2a) were significantly reduced (Fig 5B). Spread of ME was slightly reduced by ADgO(GT1a), but was increased by TNgO(GT4) (Fig 5C). Surprisingly, different effects on spread were observed for MT where TBgO(GT1c) and TNgO(GT4) reduced spread, and ADgO(GT1a) increased spread.

**Figure 5.**
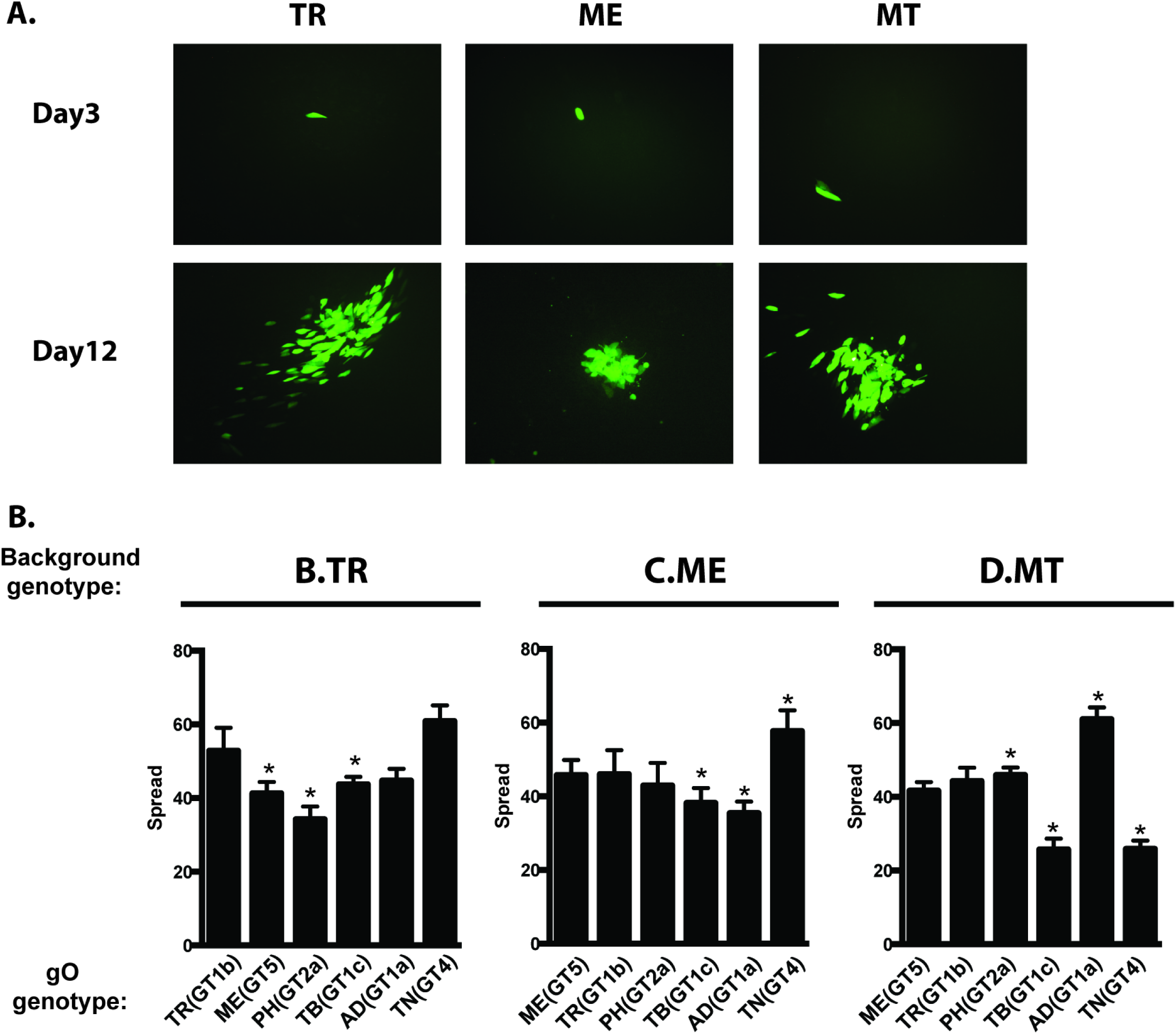
Spread of parental and heterologous gO recombinant HCMV in fibroblast cultures. Confluent monolayers of nHDF or HFFFTet (for “MT”) were infected with 0.003/cell of HCMV TR (A, B), ME (A, C), MT (A, D) or the corresponding heterologous gO recombinants. At 3 and 12 days post infection cultures were analyzed by fluorescence microscopy (A) or by flow cytometry to quantitate the total number of infected (GFP+) cells (B-D). Plotted are the average number of infected cells at day 12 per infected cell at day 3 in 3 independent experiments. Error bars represent standard deviations. Asterisks (*) denote p-values ≤ 0.05; one-way ANOVA with Dunnett’s multiple comparisons test comparing each recombinant to the parental.

A number of interesting incongruities were observed when comparing the cell-free infectivity of some gO recombinants on fibroblasts to their respective spread characteristics in fibroblasts; 1) Spread of TR_PHgO in fibroblasts was reduced compared to the parental TR (Fig 5B), but the cell-free infectivity of this recombinant was actually better (Fig 3A). Similarly, spread of both MT_TBgO(GT1c) and MT_TNgO(GT4) were reduced in fibroblasts (Fig 5D), but cell-free infectivity of both viruses was comparable to parental MT. 2) Conversely, MT_ADgO(GT1a) spread better in fibroblasts (Fig 5D), but the cell-free infectivity was substantially worse (Fig 3C). Since the efficiency of cell-free spread should depend on both the specific infectivity and the quantities of progeny virus released to the culture supernatants, it was possible that some of these incongruities reflected offsetting differences in the quantity of cell-free virus released as compared to their infectivity. To test this, progeny released from infected fibroblasts into culture supernatants were quantified by qPCR. There were no significant differences in the quantity of progeny released per cell for any of the TR or ME-based recombinants (Fig. 6A, and B). Likewise, all of MT-based recombinants released similar numbers of cell-free progeny except for MT_ADgO(GT1a), which was reduced by approximately 4-fold (Fig. 6C). Thus, the discrepancies between efficiency of spread and cell-free infectivity could not be explained by offsetting differences in the release of cell-free progeny. Rather, these results suggested that gO polymorphisms can differentially influence the mechanisms of cell-free and cell-to-cell spread in fibroblasts. The interpretation that gH/gL/gO can provide a specific function for cell-to-cell spread was supported by the results that expression of ADgO(GT1a) and TNgO(GT4), respectively reduced and increased spread of the strain ME, for which spread is almost exclusively cell-to-cell (Fig 5C).

**Figure 6.**
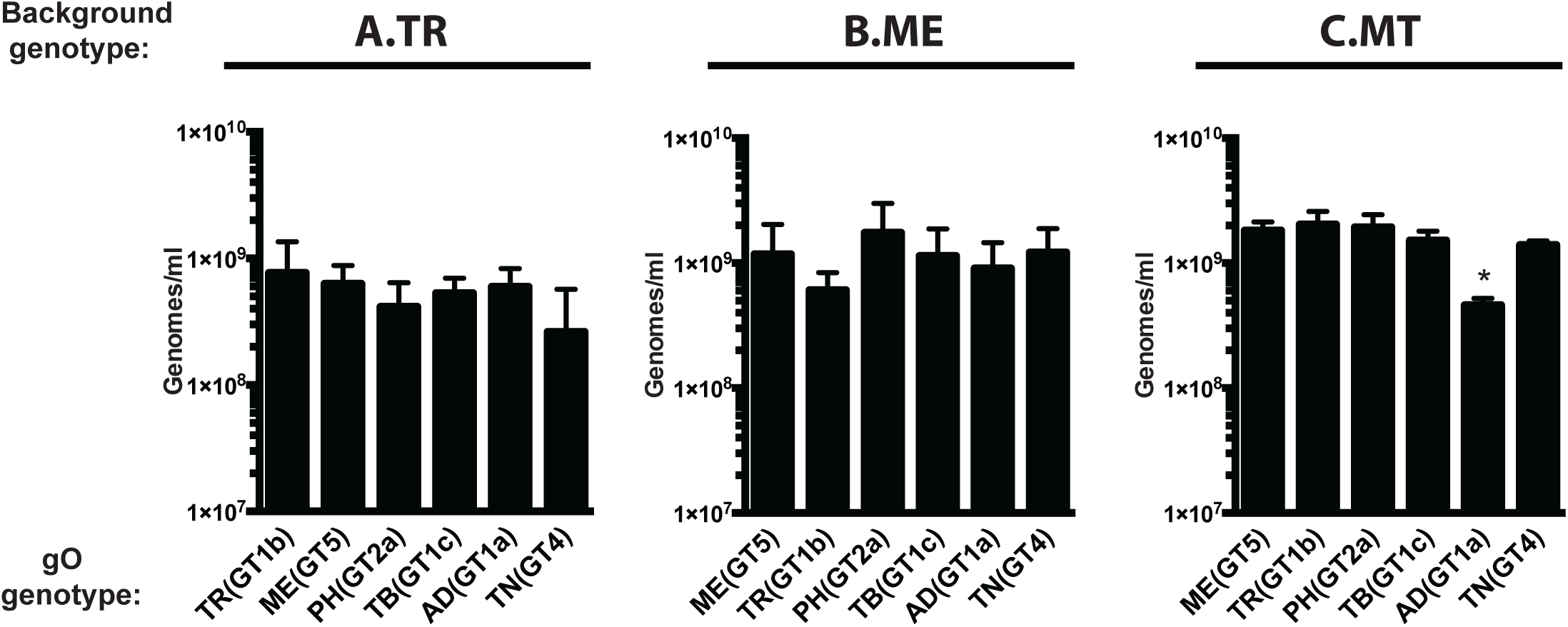
Release of extracellular progeny by parental and heterologous gO recombinant HCMV in fibroblast cultures. Cultures of nHDF or HFFFTet (for “MT”) were infected with 1 IU/cell of HCMV TR (A), ME (B), MT (C) or the corresponding heterologous gO recombinants for 8 days. The number of infected cells was determined by flow cytometry and progeny virus in culture supernatants was quantified by qPCR for viral genomes. The average number of extracellular virions per mL in each of 3 independent experiments is plotted. Error bars represent standard deviations. Asterisks (*) denote p-values ≤ 0.05; one-way ANOVA with Dunnett’s multiple comparisons test comparing each recombinant to the parental.

Spread was also analyzed in epithelial cell cultures. Here, foci of both TR and ME remained tightly localized, suggesting predominantly cell-to-cell modes of spread for both strains in this cell type (Fig. 7A). The number of TR-infected cells increased by only 5-6 fold over 12 days compared to approximately 25-fold for ME (Fig 7B and C). The low efficiency of spread for TR in epithelial cells compared to ME was documented previously and may relate to the low expression of gH/gL/UL128-131 by TR compared to ME (23, 26, 55). Expression of TNgO(GT4) further reduced TR spread in epithelial cells (Fig 7B). In contrast, ME spread was slightly reduced by TBgO(GT1c) and ADgO(GT1a), but nearly doubled by TNgO(GT4). The observed increase in ME spread due to TNgO(GT4) was not attributed to increased release of progeny to the culture supernatants in epithelial cells (Fig 8). Note that spread of MT could not be addressed in epithelial cells, since gH/gL/UL128-131 is clearly required for spread in these cells and its repression would complicate analysis of the contribution of gO polymorphisms (23). Nevertheless, it is clear from these experiments that gO polymorphisms can affect spread in epithelial cells and that this can depend on the background strain. Specifically, TNgO(GT4) reduced TR spread but increased ME spread. This suggested that although gH/gL/UL128-131 is required for efficient cell-to-cell spread in epithelial cells, and may even be sufficient in the case of gO-null HCMV (25, 27), gH/gL/gO may also contribute to the mechanism when present.

**Figure 7.**
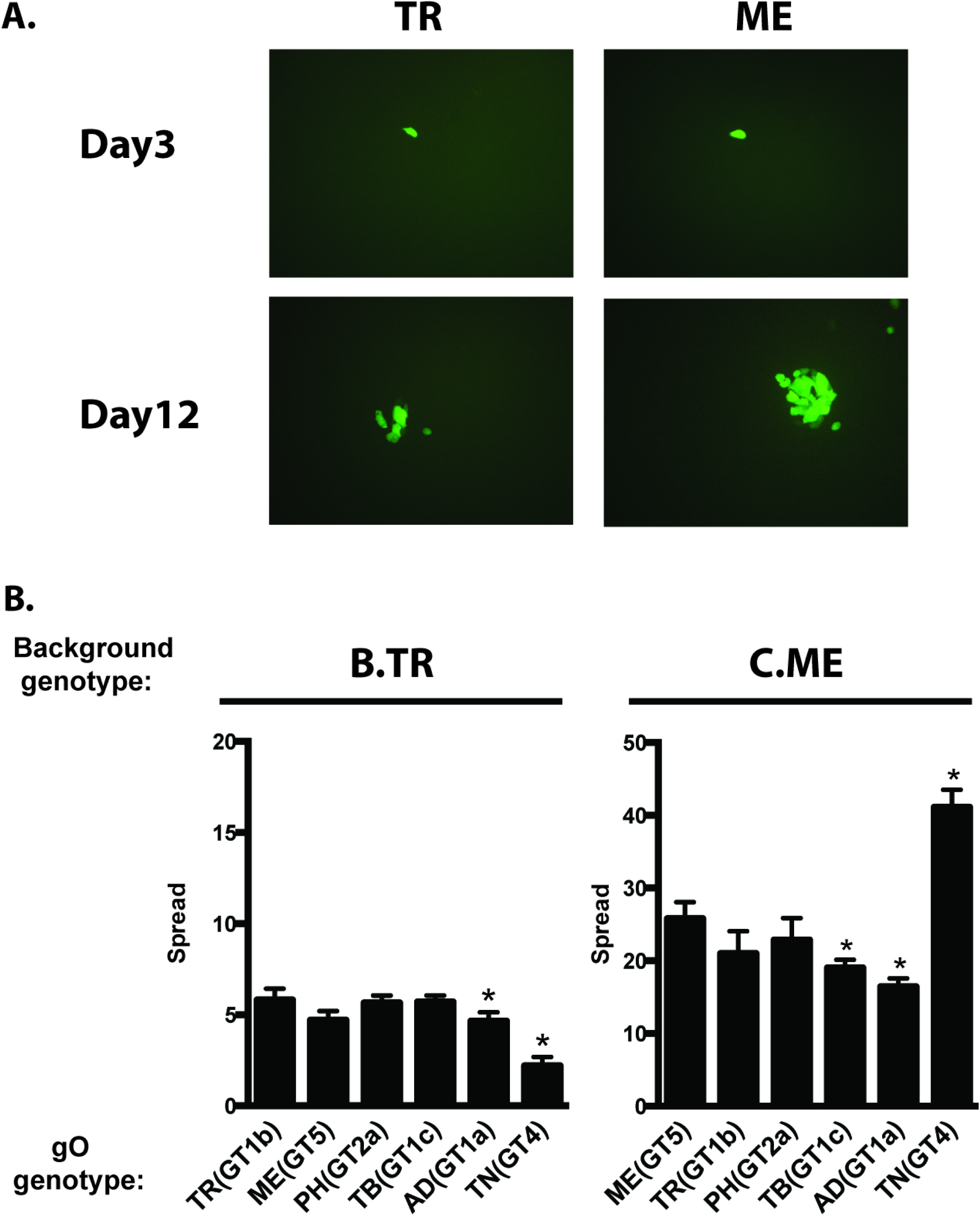
Spread of parental and heterologous gO recombinant HCMV in epithelial cell cultures. Confluent monolayers of ARPE19 cells were infected with 0.003 IU/cell of HCMV TR (A, B), ME (A, C), or the corresponding heterologous gO recombinants. At 3 and 12 days post infection cultures were analyzed by fluorescence microscopy (A) or by flow cytometry to quantitate the total number of infected (GFP+) cells (B-D). Plotted are the average number of infected cells at day 12 per infected cell at day 3 in 3 independent experiments. Error bars represent standard deviations Asterisks (*) denote p-values ≤ 0.05; one-way ANOVA with Dunnett’s multiple comparisons test comparing each recombinant to the parental.

**Figure 8.**
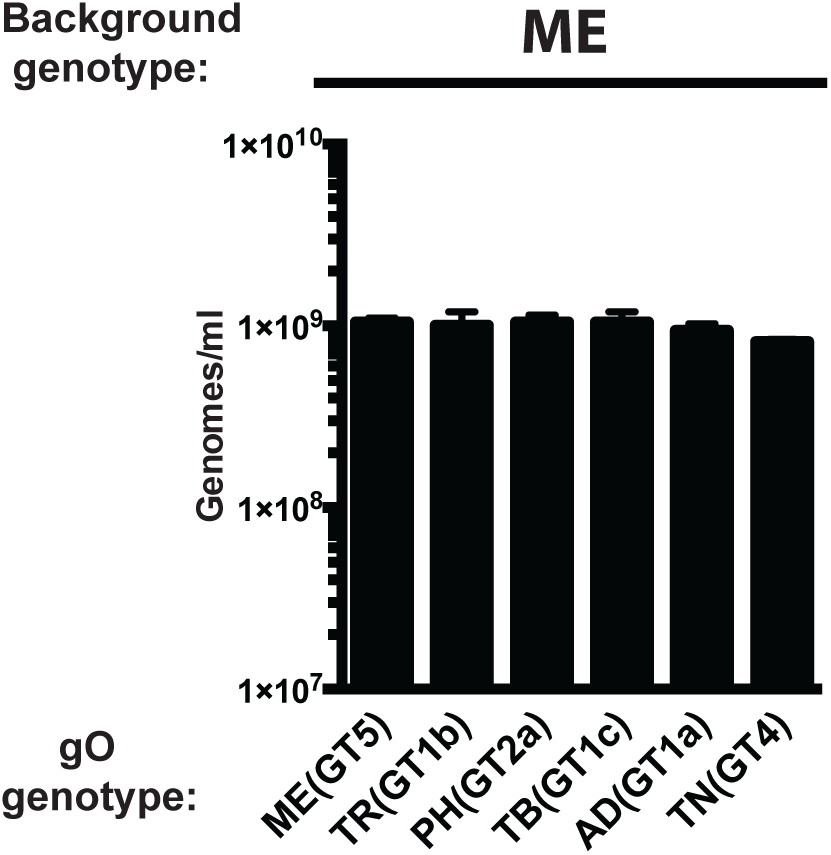
Release of extracellular progeny by parental and heterologous gO recombinant HCMV ME in epithelial cell cultures. Cultures of ARPE19 epithelial cells were infected with HFFF-tet-derived MT or corresponding heterologous gO recombinants at the highest multiplicities possible given the specific infectivity of stocks reported in Fig 3 (approximately 0.0005 IU/cell). (Note: since APRE19 cells do not express TetR, after the initial infection, MT replicates as ME). Cultures were then propagated by trypsinization and reseeding of intact cells until the number of infected cells approached 90-100% by microscopy inspection for GFP+ cells. After 8 more days, culture supernatants were then analyzed by quantified by qPCR for viral genomes. The average number of extracellular virions per mL in each of 3 independent experiments is plotted. Error bars represent standard deviations. Asterisks (*) denote p-values ≤ 0.05; one-way ANOVA with Dunnett’s multiple comparisons test comparing each recombinant to the parental.

### Polymorphisms in gO can affect antibody neutralization on gH epitopes

The extensive N-linked glycosylation of gO raised the possibility that gO could present steric hindrance to the binding of antibodies to epitopes on gH/gL, as was shown for HCMV gN and also HIV env (44, 45). A corollary hypothesis was that such effects might vary with the polymorphisms among gO isoforms. To address this, neutralization experiments were conducted using two monoclonal anti-gH antibodies; 14-4b, which recognizes a discontinuous epitope likely located near the membrane proximal ectodomain of gH (35, 56) and AP86, which binds to a continuous epitope near the N-terminus of gH (57). Note that these experiments could only be performed with TR- and MT-based recombinants since the cell-free progeny of ME-based viruses were found to be only marginally infectious (Fig 3B).

Parental TR and recombinants encoding MEgO(GT5), PHgO(GT2a) and TBgO(GT1c) were completely neutralized on fibroblasts by mAb 14-4b, whereas TR_ADgO(GT1a) and TR_TNgO(GT4) were significantly resistant (Fig 9A). There was more variability among TR-based recombinants with mAb AP86 (Fig 9B). Here, parental TR could only be neutralized to approximately 40% residual infection. TNgO(GT4) rendered TR totally resistant to mAb AP86, and MEgO(GT5) also significantly protected TR. In contrast, TR_TBgO(GT1c) and TR_ADgO(GT1a) were more sensitive to mAb AP86. On epithelial cells neutralization by both antibodies was more potent and complete than on fibroblasts, and there was less variability among gO recombinants (Fig 9C, and D). This was consistent with the interpretation that both 14-4b and AP86 could bind their epitopes on gH/gL/UL128-131 and that this represented the majority of the observed neutralization on epithelial cells. However, TR_TNgO(GT4) still displayed some reduced sensitivity to both antibodies, suggesting that gH/gL/gO epitopes also contributed to neutralization on epithelial cells.

**Figure 9.**
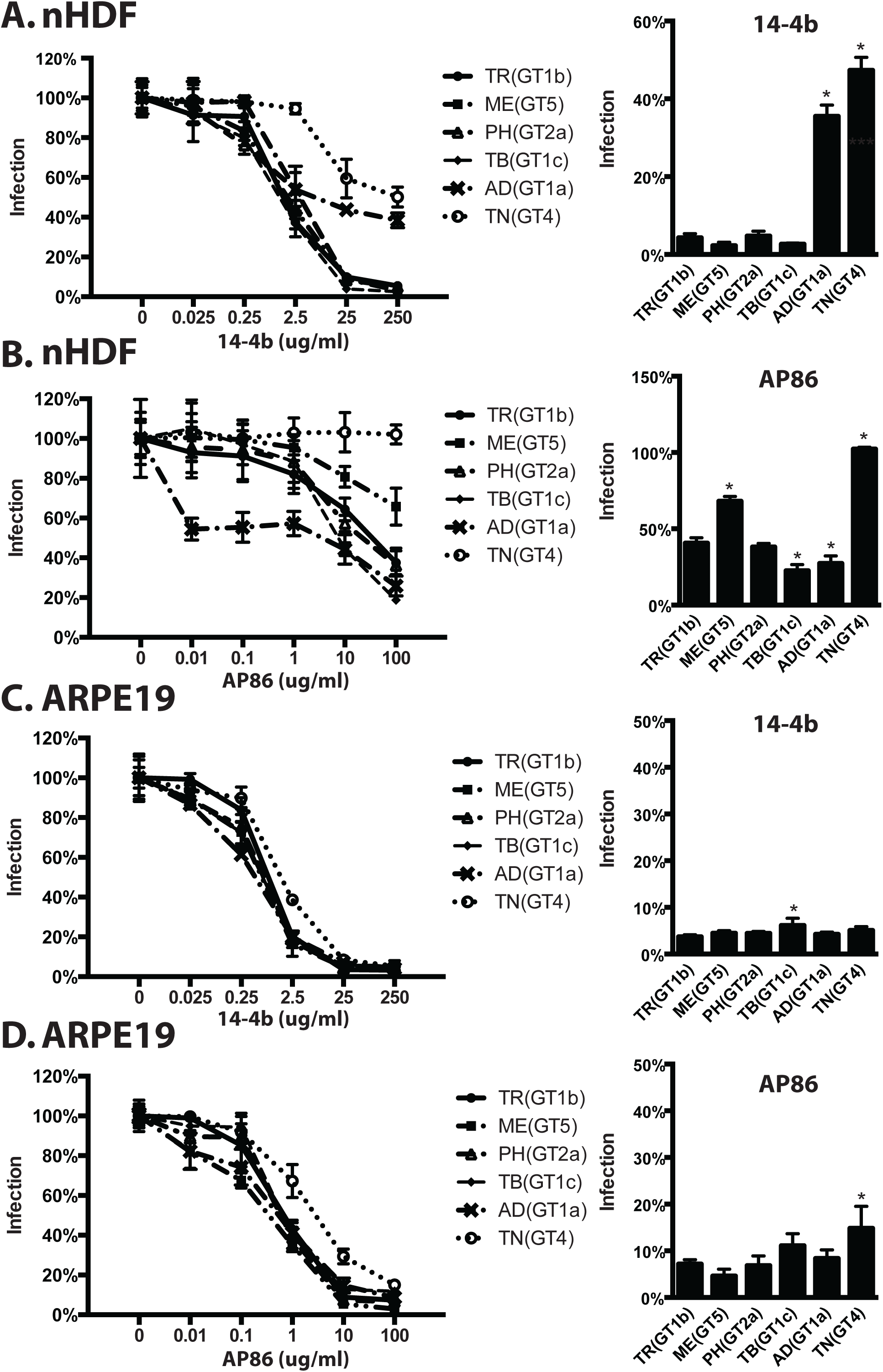
Neutralization of parental HCMV TR and heterologous gO recombinant by anti-gH antibodies. Genome equivalents of extracellular HCMV TR or the corresponding heterologous gO recombinants were incubated with 0.025-250 μg/mL of anti-gH mAb 14-4b, or 0.01-100 μg/mL of anti-gH mAb AP86 and then plated on cultures of nHDF fibroblasts (A and B) or ARPE19 epithelial cells (C and D). At 2 days post infection the number of infected (GFP+) cells was determined by flow cytometry and plotted as the percent of the no antibody control. (Left panels) Full titration curves shown are representative of three independent experiments, each performed in triplicate. (Right panels) Average percent of cells infected at the highest antibody concentrations in 3 independent experiments. Error bars represent standard deviations. Asterisks (*) denote p-values ≤ 0.05; one-way ANOVA with Dunnett’s multiple comparisons test comparing each recombinant to the parental.

MT-based recombinants were generally more sensitive to neutralization by 14-4b than were TR-based viruses (compare 14-4b concentrations in Fig 9A and 10A). Strikingly, whereas TNgO(GT4) conferred 14-4b resistance to TR, it did not in MT, and instead ADgO(GT1a) provided resistance to 14-4b (Fig 10A). As was observed for TR-based recombinants, 14-4b neutralization on epithelial cells was less affected by gO polymorphisms (Fig 10B). Note that neutralization of MT-based recombinants by AP86 could not be tested since MEgH harbors a polymorphism in the linear AP86 epitope that precludes reactivity (57). Together, these results indicated that differences among gO genotypes can differentially affect antibody neutralization on gH epitopes. Moreover, which gO genotype could protect against which antibody depended on the background strain, suggesting the combined effects of gO polymorphisms and gH/gL polymorphisms.

**Figure 10.**
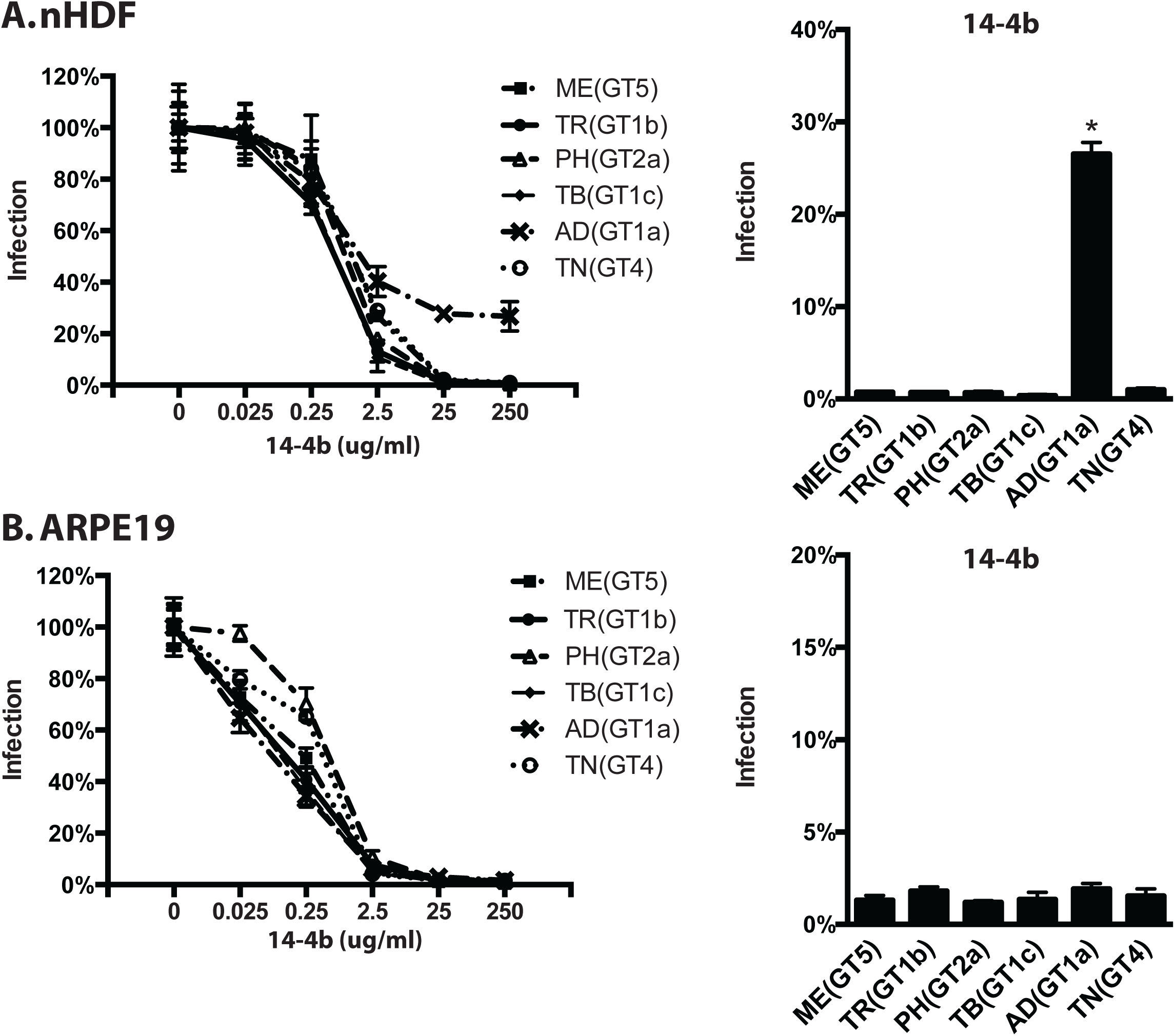
Neutralization of parental HCMV MT and heterologous gO recombinant by anti-gH antibodies. Genome equivalents of extracellular extracellular HCMV MT or the corresponding heterologous gO recombinants were incubated with 0.025-250 μg/mL of anti-gH mAb 14-4b and then plated on cultures of nHDF fibroblasts (A) or ARPE19 epithelial cells (B). At 2 days post infection the number of infected (GFP+) cells was determined by flow cytometry and plotted as the percent of the no antibody control. (Left panels) Full titration curves shown are representative of three independent experiments, each performed in triplicate. (Right panels) Average percent of cells infected at the highest antibody concentrations in 3 independent experiments. Error bars represent standard deviations. Asterisks (*) denote p-values ≤ 0.05; one-way ANOVA with Dunnett’s multiple comparisons test comparing each recombinant to the parental.

## DISCUSSION

Efficient cell-free infection of most, if not all cell types requires gH/gL/gO (22, 25, 26). However, the details of the mechanisms, and the distinctions between the roles of gH/gL/gO in cell-free and cell-to-cell spread remain to be clarified. While there are naturally occurring amino acid polymorphisms in each subunit of gH/gL/gO, gO has the most dramatic variation, with 8 known genotypes (or alleles) that differ between 10-30% of amino acids (9–12). All isoforms of gO are predicted to have extensive N-linked glycan modifications and some of the amino acid differences alter the predicted sites. In a previous report, we sought to determine if gO polymorphisms were a factor influencing the different levels of gH/gL/gO and gH/gL/UL128-131 in strains TR and ME. On the contrary, results suggested that genetic differences outside the UL74(gO) ORF result in more rapid degradation of gO in the ME-infected cells compared to TR, and this influences the pool of gO available during progeny assembly (48). Kalser et al. reported that gO polymorphisms could differentially affect multi-step replication kinetics in fibroblasts and epithelial cells (49). However, only TB was analyzed as the background and distinctions between effects on cell-free and cell-to-cell spread were unclear. In this report we constructed a matched set of heterologous gO recombinants in the well-characterized, BAC-cloned strains TR and ME. Studies included address aspects of cell-free and cell-to-cell spread, cell-type tropism and neutralization by anti-gH antibodies. The results demonstrate that gO polymorphisms can influence each of these parameters and the effects in some cases were dependent on the genetic background, suggesting a number of possible epistatic phenomena at play.

A commonly used measure to assess the tropism of HCMV strains, isolates and recombinants is the ratio of infection between fibroblasts and other cell types, including epithelial and endothelial cells (49, 55, 58, 59). Expressions of this ratio have varied, but have generally involved a normalization of the epithelial or endothelial infection to that of fibroblasts. Here we similarly determined the infectious titer of each of the parental strains and heterologous gO recombinants on both fibroblasts and epithelial cells and expressed ratios ≥1 (either fibroblasts/epithelial or epithelial/fibroblasts) to indicate the fold cell type preference or tropism of each virus (Fig 2). Both gH/gL/gO-rich viruses, TR and MT, were strongly fibroblast-tropic and some heterologous gO isoforms enhanced this preference, while others reduced it. In contrast, the gH/gL/UL128-131-rich virus ME infected both cell type more equally (ratios closer to 1), and gO polymorphisms had little effect. The limitation of any such measure of relative tropism is that it does not determine whether the virus in question can efficiently infect one cell type in particular, both or neither. Thus, any 2 viruses compared may have the same fibroblast-to-epithelial cell infectivity ratio for completely different reasons. To address this we also compared infectivity on both cell types using a common comparison for all viruses, i.e., the number of virions in the stock as determined by qPCR for DNAse-protected viral genomes in the cell-free virus stocks (Fig 3). This analysis provided a measure of specific infectivity as the number of genomes/IU, where the lower ratio indicates more efficient infection. Whether higher genomes/IU values reflect the presence of greater numbers of *bona fide* “defective” virions, or a lower probability or efficiency of each viable virion in the stock to accomplish a detectable infection, and whether or how these two possibilities are different is difficult to know for any type of virus. Nevertheless, these analyses provided important insights to the tropism ratios reported. In general, the specific infectivity ratios of the gH/gL/gO-rich viruses TR and MT in these experiments were in the range of 500-5000 genomes/IU on fibroblasts, but these viruses were approximately 20-100 fold less infectious on epithelial cells, explaining the strong fibroblast preference exhibited by these strains. The effect of most heterologous gO isoforms was similar on both cell types, but often of larger magnitude on fibroblasts. Thus, while all of the TR and MT-based gO recombinants remained fibroblast tropic, the quantitatively different effects on the two cell types influenced the magnitude of fibroblasts preference. Importantly, in no case did the change of gO affect the fundamental fibroblast preference of either TR or MT. The infectivity of the gH/gL/UL128-131-rich, ME-based viruses on both cell types was undetectable in these assays. Thus, the near neutral fibroblast-to-epithelial tropism ratios of the ME-based viruses seem to reflect an equal inability to infect either cell type and any assertion of a “preference” for either cell type for extracellular ME virions seems spurious.

Binding to PDGFRα through gO is clearly critical for infection of fibroblasts (30). However, while gH/gL/gO is also important for infection of epithelial cells, the literature is conflicted on the expression of PDGFRα and its importance for HCMV infection in epithelial and endothelial cells (26, 28, 29, 32, 33). On either cell type, possible mechanisms of gH/gL/gO include facilitating initial attachment to cells, promoting gB-mediated membrane fusion, and signaling though PDGFRα or other receptors. While Wu et al. were able to coimmunoprecipitate gB with gH/gL/gO and PDGFRα, Vanarsdall et al. showed that gH/gL without gO or UL128-131 can directly interact with gB and promote gB-fusion activity (20, 32, 34). It has also been shown that gH/gL/gO engagement of PDFGRα can elicit signaling cascades, but that this is not required for infection (28, 30, 32). In contrast, there is evidence that gH/gL/gO can help facilitate initial virion attachment (33, 54). In our studies, TNgO(GT4) reduced binding of TR to both fibroblasts and epithelial cells (Fig 4, Tables 2 and 3). However, the reduced binding of TR_TNgO(GT4) did not result in reduced infection of either cell type, and there were other isoforms of gO that either resulted in increased or decreased infectivity but were not associated with any detectable alteration in binding. Thus, while gH/gL/gO may contribute to initial binding, it is likely involved in other important mechanisms that facilitate infection and these can be influenced by gO polymorphisms. For example, it is possible that polymorphisms in gO can affect the nature and outcome of PDGFRα engagement. In support of this hypothesis, Stegmann et al. showed that mutation of conserved residues within the N-terminal variable domain of gO were critical for PDGFRα binding (60). Thus it is conceivable that the variable residues of gO can alter the architecture of the interaction with PDGFRα. Alternatively, it may be that there are other receptors on both cell types for gH/gL/gO and that gO polymorphisms can affect those interactions. Also, the effects of several specific gO isoforms observed in the TR-background were not observed in the ME or MT-backgrounds. Possible explanations for the apparent epistasis include not only the differential contributions of polymorphisms in gH/gL, but also potential differences between strains in other envelope glycoproteins, such as gB, or gM/gN may influence the relative importance of gH/gL/gO for binding and infection.

The mechanistic distinctions between cell-free and cell-to-cell spread of HCMV are unclear. Spread of ME in both fibroblast, epithelial and endothelial cells is almost exclusively cell-to-cell and this can be at least partially explained by the non-infectious nature of cell-free ME virions (Fig 3) (27, 50, 51, 55). Laib Sampaio et al. showed that inactivation of the UL74(gO)ORF in ME did not impair spread but that a dual inactivation of both gO and UL128 completely abrogated spread (27). This indicates that gH/gL/UL128-131 is sufficient for cell-to-cell spread in fibroblasts or endothelial cells in the absence of gH/gL/gO, and it seems likely that spread in epithelial cells might be similar in this respect. Our finding that various heterologous gO isoforms can enhance or reduce spread of ME without affecting the cell-free infectivity strongly suggest that while gH/gL/UL128-131 may be sufficient for cell-to-cell spread, gH/gL/gO can modulate or mediate the process, if present in sufficient amounts. In the context of MT, where expression of gH/gL/UL128-131 is reduced to sub-detectable levels (26, 51) the virus gained cell-free spread capability, and yet some of the heterologous gO isoforms had opposite effects on cell-free infectivity and spread (compare Fig 3C to 5D). Similar discorrelations between cell-free infectivity and spread were observed for the naturally gH/gL/gO-rich strain TR, albeit with different heterologous gO isoforms involved. That gO polymorphisms can have opposite effects on cell-free and cell-to-cell spread supports a hypothesis of mechanistic differences in how gH/gL/gO mediates the two processes, and again these effects seem dependent on epistatic influences of the different genetic backgrounds.

Beyond the roles of gH/gL/gO in replication, the complex is likely a significant target of neutralizing antibodies, and therefore a valid candidate for vaccine design. Several groups have reported neutralizing antibodies that react with epitopes contained on the gH/gL base of both gH/gL/UL128-131 and gH/gL/gO and others that react to gO (35–43). We found that changing the gO isoform can have dramatic effects on the sensitivity to two anti-gH mAbs (Figs 9 and 10). In the TR background on fibroblasts, both ADgO(GT1a) and TNgO(GT4) conferred significant resistance to neutralization by 14-4b, which likely reacts to a discontinuous epitope near the membrane proximal ectodomain of gH (35, 56). TNgO(GT4) also conferred resistance to AP86, which reacts to a linear epitope near the N-terminus of gH (57), whereas ADgO(GT1a) actually increased sensitivity of TR to AP86. Neutralization by either antibody on epithelial cells was not significantly affected, consistent with the notion that these antibodies can also neutralize by reacting to gH/gL/UL128-131. Again, the strain background exerted considerable influence over the effects of gO polymorphisms. For MT, it was ADgO(GT1a) that conferred resistance to 14-4b, and the other isoforms had little or no effect. The observed effects on neutralization on gH epitopes likely involve differences in how gO variable regions or associated glycans fold onto gH/gL to exert differential steric effects. Relatedly, the differential influence of gO isoforms in the two genetic backgrounds suggests epistasis involving the additive effects of gO polymorphisms with the more subtle gH polymorphisms, which together can differentially affect the global conformation of the gH/gL/gO trimer.

Previous analyses have suggested two groups of gH sequences defined by polymorphisms at the N-terminus, including the AP86 epitope (57, 61). Of the strains represented in this study, TB, TR and AD belong to the gH1 genotype and are sensitive to AP86, whereas ME, TN and PH belong to gH2 genotype and are resistant to AP86. The differential effects of gO recombinants reported here raise questions about the combinations of gH and gO genotypes in HCMV circulating in human populations. The recently published genome sequence datasets from clinical specimens have been collected with short-read sequencing approaches, which allow sensitive detection of the various gH and gO genotypes within samples, but not the combinations of the two ORFs on individual genomes (1, 3, 4, 6, 7). To address, this we analyzed 236 complete HCMV genome sequences of isolated strains and BAC clones in the NCBI database (Fig 11). Approximately half the sequences were gH1 and the other half gH2. ADgO(GT1a) and TBgO(GT1c) genotypes were exclusively linked to gH1, whereas MEgO(GT5) was exclusively linked to gH2. Other gO genotypes were found mixed with both gH genotypes, but in most cases, disproportionally with one of the gH genotypes. These analyses agreed with Rasmussen et al who suggested a strong linkage between gH1 and gO1 genotypes (note that their study predated the GT1a, 1, b, and 1c subdivisions) (10). Thus, it appears that gH and gO genotypes are non-randomly linked. This may be due in part to the adjacent position of UL74(gO) and UL75(gH) on the HCMV genome and the sequence diversity, together limiting the frequency of recombination, as suggested by the high linkage-disequilibrium of this region reported by Lassalle et al (3). In addition, our results may suggest linkage pressures based on functional compatibility of gH and gO. However, it was worth noting that among the more striking effects reported were the loss of cell-free infectivity and differential sensitivity to neutralization by gH antibodies of TR_ADgO(GT1a). Together, with the fact that TR and AD are of the same gH genotype, these results suggest epistatic interplay of genetic variation of other loci with that of gH and gO.

**Figure 11.**
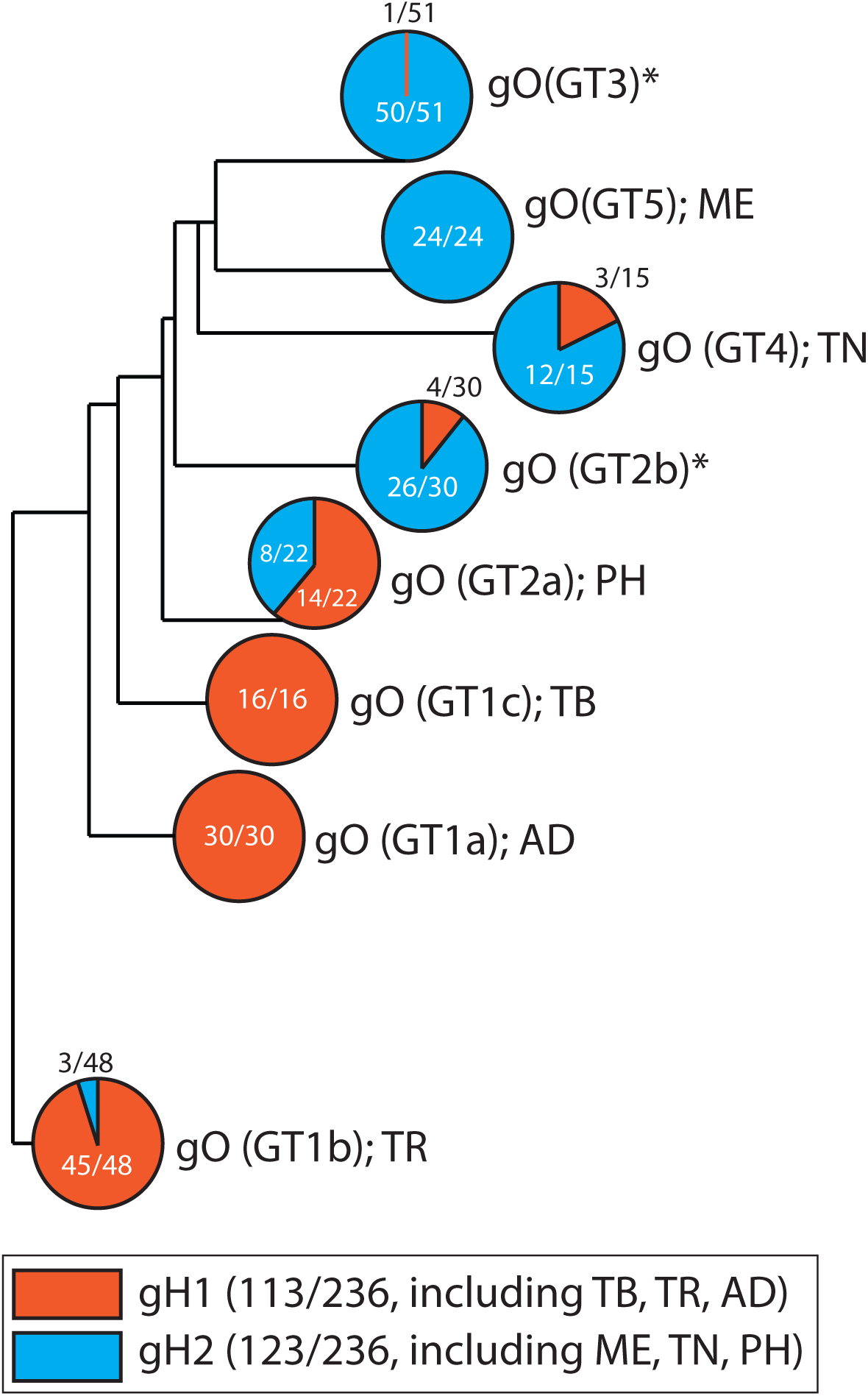
Association of gH and gO genotypes in 236 complete HCMV genome sequences in the NCBI database. Complete HCMV genome sequences were retrieved from the NCBI nucleotide database using the keywords filter human herpesvirus type 5 complete genome. The resulting set of 350 sequences was curated to remove duplicates or genomes missing any of the UL74(gO) and UL75(gH) open reading frames, generating a working set of 236 complete HCMV genomes, which were analyzed using MAFFT FFT-NS-i (v7.429) phylogeny software. UL74(gO) and UL75(gH) sequences were assigned to their respective genotype groups as defined previously; UL75(gH) genotypes 1 and 2 (57, 61); UL74(gO) genotypes 1a, 1b, 1c, 2a, 2b, 3, 4 and 5 (10, 12). Shown is a phylogenetic tree of the 8 gO genotypes with the frequency of pairing with either gH1 or gH2. Asterisks (*) indicate gO genotypes that were not analyzed in the experiments described herein.

In conclusion, we have shown that naturally occurring polymorphisms in the HCMV gO can have a dramatic influence on significant aspects of HCMV biology including, cell-free and cell-to-cell spread, and neutralization by anti-gH antibodies. These effects could not be explained by changes to the levels of gH/gL complexes in the virion envelope, but rather point to changes in the mechanism(s) of gH/gL/gO in the processes of cell-free and cell-to-cell spread. The associated epistasis with the global genetic background highlights a particular challenge for intervention approaches since humans can be superinfected with several combinations of HCMV genotypes and recombination may occur frequently (1–8). Moreover, these observations could help explain the incomplete protection observed for the natural antibody response against HCMV.

## MATERIALS AND METHODS

### Cell lines

Primary neonatal human dermal fibroblasts (nHDF; Thermo Fisher Scientific), MRC-5 fibroblasts (ATCC CCL-171; American Type Culture Collection), and HFFFtet cells (which express the tetracycline [Tet] repressor protein; provided by Richard Stanton) (51) were grown in Dulbecco’s modified Eagle’s medium (DMEM; Thermo Fisher Scientific) supplemented with 6% heat-inactivated fetal bovine serum (FBS; Rocky Mountain Biologicals, Inc., Missoula, MT, USA) and 6% bovine growth serum (BGS; Rocky Mountain Biologicals, Inc., Missoula, MT, USA) and and with penicillin streptomycin, gentamycin and amphotericin B. Retinal pigment epithelial cells (ARPE19) (American Type Culture Collection, Manassas, VA, USA) were grown in a 1:1 mixture of DMEM and Ham’s F-12 medium (DMEM:F-12)(Gibco) and supplemented with 10% FBS and with penicillin streptomycin, gentamycin and amphotericin B.

### Human Cytomegalovirus (HCMV)

All HCMV were derived from bacterial artificial chromosome (BAC) clones. The BAC clone of TR was provided by Jay Nelson (Oregon Health and Sciences University, Portland, OR, USA) (62). The BAC clone of Merlin (ME) (pAL1393), which carries tetracycline operator sequences in the transcriptional promoter of UL130 and UL131, was provided by Richard Stanton (51). All BAC clones were modified to express green fluorescent protein (GFP) by replacing the US11 ORF with the eGFP gene under the control of the murine CMV major immediate early promoter. The constitutive expression of eGFP allows the monitoring of HCMV infection early and was strain-independent. Infectious HCMV was recovered by electroporation of BAC DNA into MRC-5 fibroblasts, as described previously by Wille et al. (25) and then coculturing with nHDF or HFFFtet cells. Cell-free HCMV stocks were produced by infecting HFF or HFFFtet cells at 2 PFU per cell and harvesting culture supernatants at 8 to 10 days postinfection (when cells were still visually intact). Harvested culture supernatants were clarified by centrifugation at 1,000 X g for 15 min. Stock aliquots were stored at −80°C. Freeze-thaw cycles were avoided. Infectious unit (IU) were determined by infecting replicate cultures of nHDF or ARPE19 with serial 10-fold dilutions and using flow cytometry to count GFP positive cells at 48 hours post infection.

### Heterologous UL74(gO) recombinant HCMV

A modified, three step BAC En Passant recombineering technique was performed (63, 64). In the first step, the endogenous UL74 ORF from the start codon to the stop codon of both TR and ME was replaced by a selectable marker. This necessary step was added to prevent formation of chimeric UL74 gene by internal recombination of the UL74 BAC sequence and the incoming heterologous UL74 ORF. A purified PCR product containing the ampicillin resistance selectable marker (AmpR) cassette from the pUC18 plasmid flanked by sequences homologous to 50 bp upstream and downstream of the TR or ME UL74 ORF was electroporated into the bacteria, recombination was induced and the recombinant-positive bacteria were selected on medium containing ampicillin (50 µg/ml) and chloramphenicol (12.5 μg/ml). The primers used to produce the TR- and ME-specific AmpR PCR bands are For74TRamp, 5’-CATGGGAGCTTTTTGTATCGTATTACGACATTGCTGTTTCCAGAACTTTAcgcggaacccctatttgtttatttttctaaatac, For74MEamp, 5’-GATGGGAGCTTTTTGTATCGTATTACGACATTGCTGCTTCCAGAACTTTAcgcggaacccctatttgtttatttttctaaatac, and Rev74amp (used for both TR and ME PCR reactions), 5’-CCAAACCACAAGGCAGACGGACGGTGCGGGGTCTCCTCCTCTGTCATGGGGttaccaatgcttaatcagtgaggcacc. The lower case nucleotides correspond to the AmpR gene from the pUC18 plasmid, the upper case nucleotides to the TR and ME BAC sequences immediately upstream and downstream of the UL74 ORF.

In the second step, the AmpR cassette in the TR and ME first-step intermediate BACs was replaced with the UL74(gO) sequence from the heterologous strain containing the En Passant cassette (63, 64). Briefly, E. coli cultures were prepared for recombination as described above for step 1 and electroporated with purified PCR products containing the UL74 ORF from the TR or ME strain flanked by sequence homologous to 50 bp upstream and downstream of the opposite strain. The UL74 ORF also contained an inserted En Passant cassette (an I-SceI site followed by a kanamycin resistance gene surrounded by a 50-bp duplication of the UL74 nucleotides of the insertion site). Transformed E. coli cells were induced for recombination and then selected for the swap of the UL74 En Passant sequence into the BAC by growth on medium containing kanamycin (50 µg/ml) and chloramphenicol (12.5 μg/ml). A PCR reaction analysis with primers located upstream and downstream of UL74 was used to confirm the swap of the AmpR cassette by the En Passant cassette/UL74 gene.

In the third step, several sequencing validated colonies of the second step were subjected to the last step of the En Passant recombineering, that is, an induction of both the I-SceI endonuclease and the recombinase (63, 64). The activity of these enzymes lead to an intramolecular recombination in the UL74 sequence around the En Passant cassette and thus the restoration of an uninterrupted, full length UL74 ORF. The final heterologous UL74(gO) recombinants were verified by Sanger sequencing of PCR products using primers located upstream and downstream of the UL74 gene.

### Antibodies

Monoclonal antibodies (MAbs) specific to HCMV major capsid protein (MCP), pp150, and gH (14-4b and AP86) were provided by Bill Britt (University of Alabama, Birmingham, AL) (35, 57, 65, 66). 14-4b and AP86 were purified by FPLC and quantified by the University of Montana Integrated Structural Biology Core Facility. Rabbit polyclonal sera against HCMV gL was described previously (9, 26).

### Immunoblotting

HCMV cell-free virions were solubilized in 2% SDS–20 mM Tris-buffered saline (TBS) (pH 6.8). Insoluble material was cleared by centrifugation at 16,000 X g for 15min, and extracts were then boiled for 10 min. For reducing blots, dithiothreitol (DTT) was added to extracts to a final concentration of 25 mM. After separation by SDS-PAGE, proteins were transferred onto polyvinylidene difluoride (PVDF) membranes (Whatman) in a buffer containing 10 mM NaHCO_3_ and 3mM Na_2_CO_3_ (pH 9.9) plus 10% methanol. Transferred proteins were probed with MAbs or rabbit polyclonal antibodies, anti-rabbit or anti-mouse secondary antibodies conjugated with horseradish peroxidase (Sigma-Aldrich), and Pierce ECL-Western blotting substrate (Thermo Fisher Scientific). Chemiluminescence was detected using a Bio-Rad ChemiDoc MP imaging system. Band densities were quantified using BioRad Image Lab v 5.1.

### Quantitative PCR

Viral genomes were determined as described previously (26). Briefly, cell-free HCMV stocks were treated with DNase I before extraction of viral genomic DNA (PureLink viral RNA/DNA minikit; Life Technologies/Thermo Fisher Scientific). Primers specific for sequences within UL83 were used with the MyiQ real-time PCR detection system (Bio-Rad).

### Flow cytometry

Recombinant GFP-expressing HCMV-infected cells were washed twice with PBS and lifted with trypsin. Trypsin was quenched with DMEM containing 10% FBS and cells were collected at 500 g for 5 min at RT. Cells were fixed in PBS containing 2% paraformaldehyde for 10 min at RT, then washed and resuspended in PBS. Samples were analyzed using an AttuneNxT flow cytometer. Cells were identified using FSC-A and SSC-A, and single cells were gated using FSC-W and FSC-H. BL-1 laser (488nm) was used to identify GFP+ cells, and only cells with median GFP intensities 10-fold above background were considered positive.

### Virus particle binding

nHDF or ARPE19 cells were seeded at density of 35,000 cells per cm^2^ on chamber slides (Nunc Lab Tek II). 2 days later, virus stocks were diluted with media to equal numbers of virus particles based on genome quantification by qPCR. Binding of virus particles to the cells was allowed for 20min at 37°C. Then the inoculum was removed, and the cells were washed once with medium to remove unbound virus before fixation and permeabilization with 80% acetone for 5min. Bound virus particles were stained with an antibody against the capsid-associated tegument protein pp150 (65) which allowed to detect enveloped particles attached to the plasma membrane as well as internalized particles. For visualization, a goat anti-mouse Alexa Fluor 488 (Invitrogen) secondary antibody was used. Unbound secondary antibody was washed off before the chambers were removed and the cells were mounted with medium containing DAPI (Fluoroshield) and sealed with a cover slide for later immunofluorescence analysis. Images were taken with a Leica DM5500 at 630-fold magnification. For each sample 10 images with 4 to 6 cells per image were taken and the number of cell nuclei as well as the number of virus particles was determined using Image J Fiji software (v 1.0). Three independent virus stocks were tested in 3 independent experiments.

### Antibody neutralization assays

Equal numbers of nHDF-derived cell-free parental viruses and heterologous gO recombinants were incubated with multiple concentrations of anti-gH mAb 14-4b or AP86 for 1hr at RT then plated on nHDF or ARPE19 for 4hrs at 37°C. Cells were then cultured in the appropriate growth medium supplemented with 2% FBS. After 2 days, cells were detected from the dish and fixed for flow cytometry analyses. Each antibody concentration was performed in triplicate and 3 independent experiments were conducted.

## ACKNOWLEDGMENTS

We are grateful to Bill Britt, David Johnson, Jay Nelson, and Richard Stanton for generously supplying HCMV BAC clones, antibodies, and cell lines as indicated in the Material and Methods, and members of the Ryckman laboratory for support, and insightful discussions. We also thank Ekaterina Voronina and Mary Ellenbecker of University of Montana for assistance with immunofluorescent microscopy, the staff of the University of Montana Center for Biomolecular Structure and Dynamics Integrated Structural Biology Core Facility for help purifying monoclonal antibodies, and the staff of the University of Montana Flow Cytometry Core of the Center for Environmental Heath Sciences for assistance with flow cytometry.

This work was supported by grant from the National Institutes of Health to B.J.R (R01AI097274), a fellowship from the German Research Foundation (DFG) to C.S. (STE 2835/1-1), a fellowship from American Heart Association to E.P.S (17POST33350043) and a National Institutes of Health CoBRE award to Center for Biomolecular Structure and Dynamics at University of Montana (PG20GM103546).

Experiments were designed by B.J.R., L.Z.D, C.S., and E.P.S, and performed by L.Z. and C.S. Critical reagents were developed by L.Z.D., J.M.L. and Q.Y. Data were analyzed, and manuscript was prepared by B.J.R., L.Z.D., C.S., Q.Y., E.P.S., and J.M.L.

